# Soft tissue constraints on joint mobility in the avian shoulder

**DOI:** 10.1101/2025.05.16.654461

**Authors:** Oliver E. Demuth, John R. Hutchinson, Vittorio La Barbera, Sharon E. Warner, Daniel J. Field

**Affiliations:** Clare College, University of Cambridge, Cambridge, UK; Department of Earth Sciences, University of Cambridge, Cambridge, UK; Structure and Motion Laboratory, Royal Veterinary College, Hatfield, UK; Museum of Zoology, University of Cambridge, Cambridge, UK

**Keywords:** Range of motion, joint mobility, shoulder, soft tissue constraints, bird, ligament, flight, XROMM, wing

## Abstract

Joints and their surrounding soft tissues facilitate and restrict vertebrate skeletal motion. Measures of maximal joint mobility provide insight into articular function and its limits on potential joint motion and thereby behaviour. In extinct vertebrates the reconstruction of joint mobility permits us to decipher shifts in locomotor evolution. Such measurements are generally limited to studies of osteological joint mobility. However, only a subset of osteologically feasible poses are biologically feasible because true joint mobility is limited by soft tissues, such as ligaments, that seldomly preserve in the fossil record. To address this issue, we implemented an *in silico* model to simulate avian glenohumeral (shoulder) movement and the constraints imposed by six ligaments on its joint mobility. We evaluated our *in silico* model of the partridge shoulder joint with measured *ex vivo* shoulder mobility using X-ray Reconstruction of Moving Morphology (XROMM). Our results indicate that modelling ligamentous constraints is integral to accurately quantifying shoulder function due to the role of ligaments in maintaining articular contact during complex glenohumeral motion. Our approach enables more confident estimates of functional joint mobility in both extant and extinct vertebrates and thereby stands to improve inferences of behaviour and musculoskeletal function in the vertebrate fossil record.

## 1. Introduction

Studying the functional and articular morphology of joints is crucial for understanding the function of vertebrate locomotor systems. Attempts to reconstruct major evolutionary transformations in locomotor evolution, such as the water-to-land transition among stem tetrapods [1,2] or the origin of avian flight [3,4] rely on accurate inferences of how extinct vertebrates moved. The articular morphologies of bones may hint at the potential mobility of a joint and how bones move relative to each other in living vertebrates [5]; however, inferences relying solely on these articular morphologies are limited if they ignore the important influence of soft tissues on joint mobility [6–10].

The mobility of a joint, that is, its range of motion (ROM) and its three-dimensional (3D) envelope, can be estimated by evaluating which joint poses are possible and which are prevented either by bony stops or by joint disarticulation. Joint mobility has often been considered an important foundation for reconstructing locomotor capabilities in extinct tetrapods [1,11–16], with mobility estimates traditionally relying on physical or digital manipulation of individual bones [17,18]. However, the last decade has seen substantial advances in comparative locomotor biomechanics with the introduction of new digital methods [19]. These digital toolsets have been adapted from the animation and 3D industry and include novel modelling approaches for musculature [20,21], reconstruction of deformed specimens and removal of taphonomic artefacts [21–24], animation and tracking of motion data [11,25–30] and estimation and systematic quantification of joint mobility [10,13,31–37]. Such standardised approaches are widely used to systematically investigate joint mobility across all three rotational degrees of freedom (DOF; i.e., rotation around the flexion/extension, abduction/adduction and long axes) and their complex interplay [37–39]. These approaches are, therefore, an important tool to infer joint function and exclude potential behaviours that would require impossible joint poses [1,5,11–15,31,36,40–44] under the assumption that these mobility estimates are generally reliable (but see [33]), and that only a subset of these poses are/were used in life (e.g., [15,31,39,45–47]). Resultant joint mobility estimates usually form the foundation upon which other parameters, such as behaviour, muscle moment arms or energy expenditure [11,34,48–53] are investigated. Erroneous exclusions of viable or the inadvertent inclusion of physiological inviable poses may, therefore, have repercussions for any downstream analyses, such as the estimation of musculotendon parameters in biomechanical models based upon joint excursion [53–55], and/or can lead to erroneous conclusions [5,33,56].

Recent studies have suggested that simulations which only incorporate three rotational DOFs may not sufficiently capture the true *ex vivo* mobility of a given joint, and additional translational DOFs may, therefore, be necessary to incorporate into simulations of joint mobility [13,33,36,56–58]. Yet, an unconstrained inclusion of joint translations to ROM simulations may lead to an extreme expansion of the estimated mobility beyond its true boundaries. Therefore, such simulations may no longer be functionally informative without the incorporation of additional constraints on joint motion, such as cartilage, muscles and/or ligaments [7,10,31,34,36,46].

Previous attempts to constrain joint mobility estimates have incorporated ligamentous constraints using the dynamic *nCloth* for simulations in Autodesk Maya [31,36] (Autodesk Inc., San Rafael, California, USA). Ligaments were simulated as a dynamic polygon surface, represented by the *nCloth*, wrapping around the bone surfaces from origin to insertion. The length of a ligament was then measured along the midline of this dynamic polygon surface [31,59]. However, these measurements, and thus the resulting soft tissue constraints, suffered from motion-dependency issues, because the dynamic *nCloth* object is inherently motion dependent. The calculated length of such a ligament is, therefore, highly dependent on the previous timestep, which may cause vastly different length estimates for an identical joint pose depending on different motion trajectories that lead to that pose, necessitating extensive sensitivity analyses [36]. This uncertainty may be further amplified if the motion (for which the ligaments are simulated) is non-sequential and includes large jumps in joint angles (e.g., if it is not directly derived from smooth experimental motion data, but based on simulations of joint mobility [12,31,36]).

Here, we present a solution to these challenges via implementation of a computationally fast approaches for both dynamic, contact-based ROM simulations and motion-independent ligamentous constraints for the study of vertebrate joint function. Our method is integrated within an existing pipeline for joint mobility analyses with the goal of improving joint mobility estimates from functional simulations of extant and extinct vertebrates.

As a case study, we apply our approach to investigate ROM in the avian shoulder. Specifically, we examine unique glenoid of extant birds [60] using *in silico* simulations in conjunction with *ex vivo* data collected through X-ray Reconstruction of Moving Morphology (XROMM) [25,26]. Previous work on birds and their ancestors has mainly focused on the hindlimb and terrestrial locomotion [31,33,39,61–63], with others only investigating specific behaviours in the forelimbs (e.g., [64–67]). However, clarifying overall joint mobility within the avian shoulder will be pivotal for improving inferences of wing function during the theropod-to-bird transition, which witnessed the evolutionary origin of avian powered flying ability—one of the most dramatic locomotor transitions in the evolutionary history of vertebrates.

## 2. Material and Methods

### 2.1 Study species

Five carcasses of Red-legged Partridges (*Alectoris rufa*; henceforth abbreviated as RLP) were obtained from a local gamekeeper (Table 1). Four specimens were selected for marker placement and experimental data collection (RLPs1-4) and an additional specimen (RLP5) was partially dissected to expose ligaments within the shoulder joint for measurement.

**Table 1.**
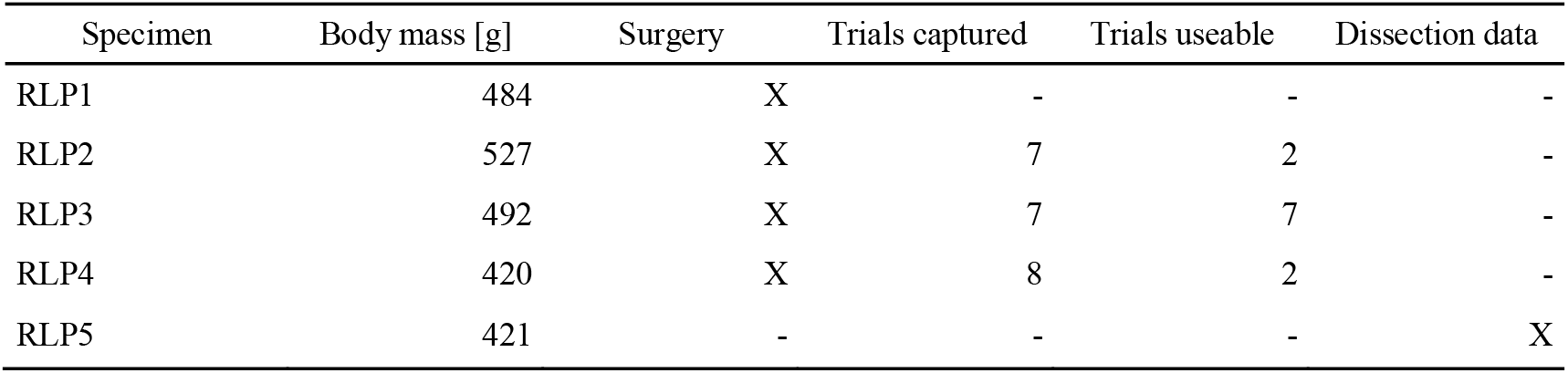
Red-legged Partridge data collection.

Sex and approximate age at time of death of the individual specimens was not known, but the birds were considered adults.

### 2.2 Marker placement and dissection

To ensure precise 3D measurements of joint movement, we implemented radio-opaque markers directly into bones [26,30,34,37,39,61,68]. Before marker implantation the specimens were visually inspected to ensure that no effects of desiccation or decomposition were evident, and to assess potential damage to the wings. Small incisions into the skin of four partridge specimens (RLP1-4) were made to expose the surface of the major bones of the pectoral girdle and wing. Subsequently, 1.1 mm holes were drilled into the exposed bones using a small hand-held drill, into which the radio-opaque 1.0 mm tantalum beads (Bal-Tec, Los Angeles, USA) were placed and fixed with cyanoacrylate glue, which was also used to close the incisions. In total 20 beads were implanted into the pectoral girdle and the bones of one wing for each specimen. Each long bone in the wing (humerus, ulna, radius and carpometacarpus), as well as the scapula and sternum, had rigid bodies defined by three tantalum beads (Figure S3). Unfortunately, due to the large amount of musculature and other soft tissue surrounding the coracoids it was not possible to implant the minimum number of markers required to accurately track their position [68]. Therefore, the position and orientation of the coracoids were approximated for the tracking (see [19,30]). Due to potential coracosternal movement (see [64]) only a dynamic plane could be defined, based on the left and right coracoid markers and the cranial sternum marker (Figure S3), rather than a rigid body.

During dissection of RLP5, the origins and insertions of six shoulder ligaments were identified (Figure 1) and verified with reference to published literature [69–71]. The lengths of each of these ligaments were measured using digital callipers sensitive to 0.1mm.

**Figure 1.**
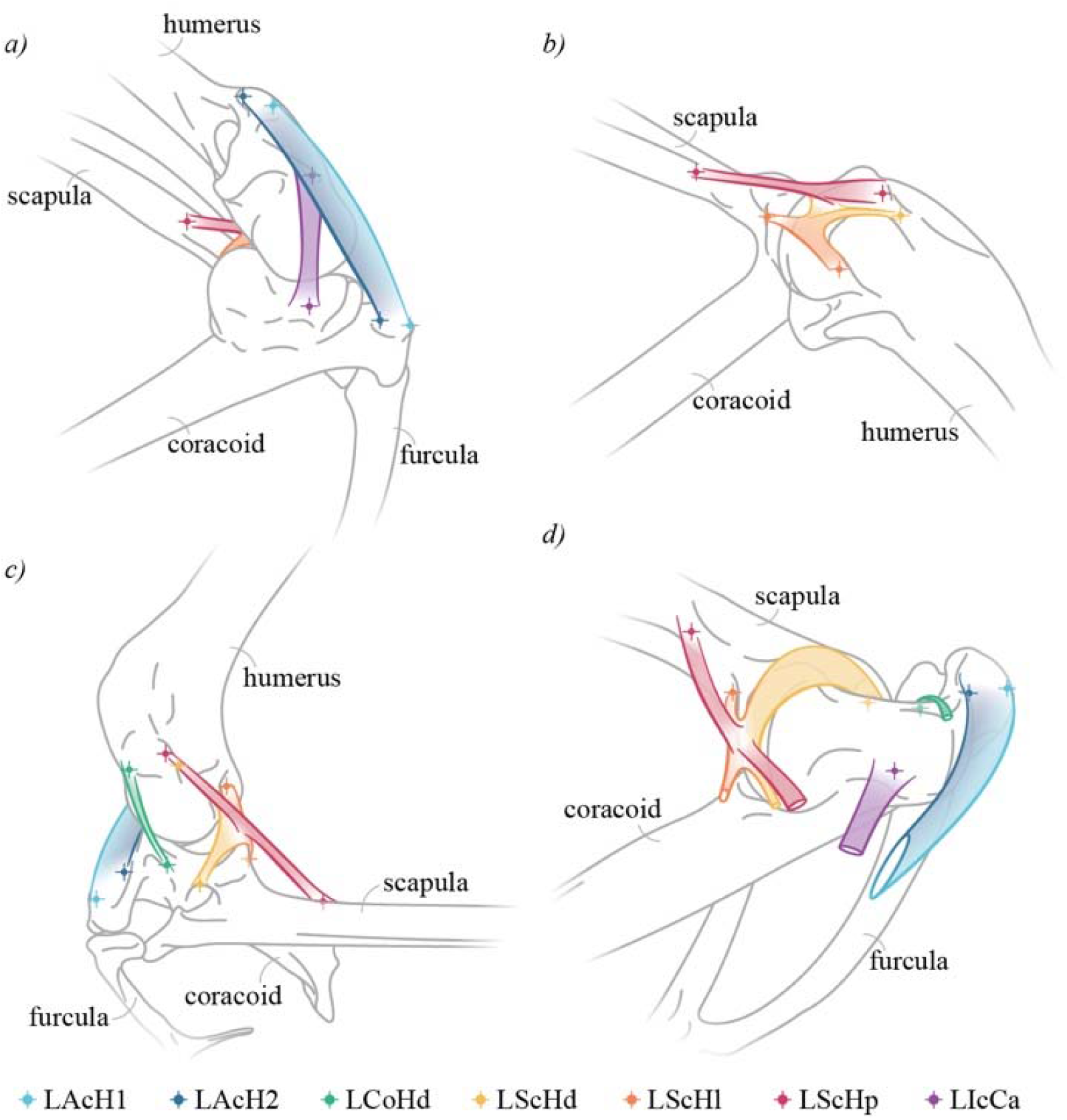
Partridge shoulder ligaments. Ligaments identified during the dissection of the shoulder capsule of the Red-legged Partridge (*Alectoris rufa*). Right shoulder joint in the following views: craniolateral (a), caudolateral (b), dorsal (c) and oblique lateral with removed humerus and cut ligaments (d). Musculature and other soft tissues have been omitted from the illustration to highlight the ligaments and the positions of their attachment sites on the scapula, coracoid, and humerus. Due to the broad attachment surfaces on the humerus and coracoid, the LAcH was divided into two separate units, 1 and 2. Crosses denote the positions of locators for ligament origins and insertions in Autodesk Maya for subsequent ligament simulations. Note the fusion of the LScH ligaments. Nomenclature follows Fürbringer [69]. Abbreviations: LAcH, *Ligamentum (L) acrocoraco-humerale*; LCoHd, *L. coraco-humerale dorsale*; LScHd, *L. scapulo-humerale dorsale*; LScHl, *L. scapulo-humerale laterale*; LScHp, *L. scapulo-humerale posterius*; LIcCa, *L. intracapsulare coracoideus anterius*.

### 2.3 Ex vivo *XROMM data collection*

*Ex vivo* movement data of the shoulders of RLPs2-4 were captured via biplanar fluoroscopy using XROMM [25,26,68]. Damage to the wings of RLP1 was identified in the μCT scan and the specimen was discarded prior to conducting experiments. The remaining partridges were placed on top of a foam board and strapped into a custom-built Velcro harness to secure their position in the capture area of the fluoroscopes. The carpometacarpus/wing was attached to a wooden manipulator rod using Velcro strips (Figure S1). The Velcro harnesses on both the body and wings and the rod itself allowed for some flexibility to reduce the risk of injuries to the joint capsules due to potentially excessive movement of the manipulator rod while attempting to achieve maximal excursion of the wing.

Two BV Libra C-arm systems (Koninklijke Philips N.V., Amsterdam, Netherlands) were used, each composed of a BV 300 generator, F017 tube and BV 300 collimator and intensifier (22.9 cm diameter), with a source-to-image distance of 99.5 cm. The X-rays were generated at 67 kV and 2.27 mA and fired continuously for the duration of a trial. The trials were recorded with two Photron FASTCAM Mini WX50 high-speed digital video cameras (Photron, Tokyo, Japan) at 250 frames per second with a resolution of 2048 × 2048 pixels and a shutter speed of 750 Hz. In total 22 trials were recorded approximating eight minutes of motion data (Table 1). X-ray videos were undistorted, calibrated and digitised using XMAL_AB_ 2.1.0 [27]. The 3D point data of the markers were filtered at 10 Hz prior to export.

### 2.4 Skeletal geometry acquisition and anatomical and joint coordinate system setup

The specimens were μCT scanned at 190 kV and 200 μA, with a voxel size of 0.125mm, at the Cambridge Biotomography Centre, Cambridge, United Kingdom, using a Nikon 49 Metrology XT H 225 ST scanner to capture both the relative marker positions and the bone geometries. μCT data were segmented using Avizo Lite 2020.2 (Thermo Fisher Scientific, Inc., Waltham, Massachusetts, USA) and the resulting meshes prepared for data analysis and simulations using MeshLab 2021 [24,72].

Anatomical coordinate systems (ACS) and joint coordinate systems (JCS) were created following established primitive shape-fitting protocols [12,53,73]. The articular surfaces of the glenoid and wing bones were first isolated in Autodesk Maya and primitive shapes were fitted in Matlab R2023b [53] (The MathWorks, Inc., Natick, Massachusetts, USA; See Supplementary Information for detailed description of the ACS setup). Based on the ACSs, the wing was collapsed into a reference pose (Figure S2) from which all translational and rotational deviations were measured [25,53,61,64,73,74].

### 2.5 In silico *ROM simulations and ROM mapping*

Range of motion (i.e., joint mobility) simulations were set up in Autodesk Maya 2025 using a novel Python implementation of the approach originally outlined by Marai and colleagues [75] and subsequently implemented in Matlab by Lee and colleagues [16], adapted for Autodesk Maya to incorporate it into existing ROM pipelines [31,32,35,47]. Our implementation builds on the theoretical framework of previous joint sampling techniques [31,32,35,47], yet has significant advantages over these approaches. Importantly, it no longer assumes a fixed joint centre position, and joint translations are optimised for each rotational joint pose. As such, it represents physiological joint movement more accurately by enabling continuous contact of articular surfaces and constant joint spacing, thus overcoming previous methodological constraints. Additionally, our simulations exploit the signed distance field (SDF) representation of the bones (i.e., as a 3D scalar field specifying the signed distance from any given field point to the bone surface [16,75,76]). This only needs to be calculated once, rather than continuously relying on computationally expensive 3D meshes; as such, our method overcomes previous computational constraints, substantially improving simulation speed.

SDFs were initialised as a cubic grid centred around the centroids of the fitted shapes to their respective articular surfaces. The signed distances were then sampled at regular intervals over the 3D grids (n = 101 x 101 x 101 grid points each). For each grid point *P*, the distance *d*_*b*_ to the bone mesh was calculated by finding the closest point *Q* on the surface of the mesh. The sign, distinguishing whether the point *P* was inside (negative) or outside (positive) a bone mesh, was determined by calculating the dot product between the vector 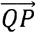 and the surface normal 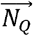 at point *Q*. If the dot product was negative, then was inside the mesh and thus its distance negative:

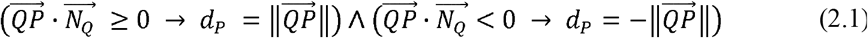

The exact signed distance to the bone surface was, therefore, known only at each grid point. The distance to the bone surface between grid points was interpolated on-demand from the surrounding grid points using tricubic interpolation [77]:

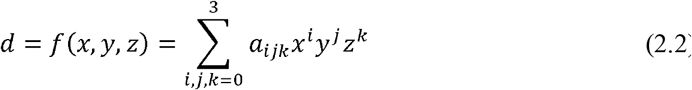

where *a*_*ijk*_ are the coefficients for each of the eight corner points of a grid node [77] and *x, y* and *z* are the Cartesian 3D coordinates of the sampled point.

We implemented a multiterm objective function *C*_*ROM*_ to optimise the translational joint position for each given joint orientation:

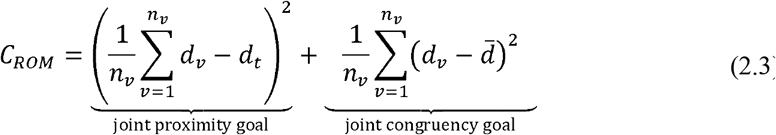

subject to

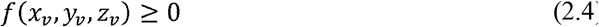

where *d*_*v*_ is the calculated signed distance for vertex *v, n*_*V*_ is the number of vertices of the articular surfaces, *d*_*t*_ is the target joint proximity and 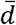 is the mean joint proximity, following [16]. Additionally minimising the variance of the joint proximity ensures maximal congruence between the articular surfaces of the proximal and distal bones, especially in non-spherical joints, such as the particularly open hemi-sellar avian shoulder joint [36,60,78]. This optimisation problem was solved using a sequential quadratic programming method (SQP) [79] and implemented in Python 3.11 for Autodesk Maya 2025 using the Python modules NumPy 1.24.4 [80], SciPy 1.15.0 [81] and pytricubic 1.0.4 [82]; see Supplementary Information.

Joint orientations were sampled across the flexion/extension (FE; −180° to 180°), abduction/adduction (ABAD; −90° to 90°) and long-axis (LAR; −180° to 180°) axes at 5° intervals, resulting in 197,173 tested joint orientations per specimen. As the joint centre position was optimised for each rotational pose, the resulting simulation inherently described six DOFs with an optimal joint translation for any given rotational pose, while simultaneously avoiding potentially excessive disarticulations of the joint [33,35].

### 2.6 Motion-independent soft tissue simulations

Furthermore, we implemented a motion-independent solution for soft tissue simulations into the existing pipeline of ROM simulations in Autodesk Maya, to circumvent motion-dependency issues related to the dynamic *nCloth* object. To do so, the path minimising algorithm of Marai and colleagues [76] was implemented in Python for Autodesk Maya 2025 to calculate the length of a ligament at any given joint pose independent of the previous time step. The 3D geometries of each bone were again approximated as SDFs. The ligament path and length could then be optimised over the much faster interpolated scalar field representation of the bones rather than relying on their computationally expensive 3D meshes [76].

Individual ligament coordinate systems were defined to facilitate the calculation of the wrapping around the bones. A vector from the origin *p*_0_ to insertion *p*_*n*_ represented the *X*-axis, which was subsequently subdivided into (n = 20) equidistant segments between *p*_0_ and *p*_n_. The *Y*-axis was defined by the cross product of the *X*-axis and a vector from the origin *p*_0_ to the joint centre *p*_*j*_, with the *Z*-axis perpendicular to both *X*- and *Y*-axes. The ligament path could then be formulated as the following optimisation problem: Find the coordinates of the *n* − 1 points, between *p*_0_ and *p*_*n*_, so that the Euclidean distance of the path along *p*_0_, *p*_1_,*p*_2_, …, *p*_*n*_ is minimal while the distance between each path point and the bony obstacles is non-negative. Because the points were equally spaced along the *X*-axis (i.e., their distance was constant) the length of the shortest path could be calculated by minimising its Euclidean distance over only the and coordinates for each point [76], which we implemented as the following cost function:

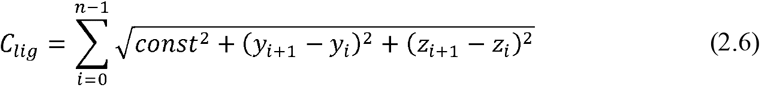

subject to

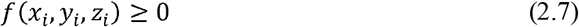

and

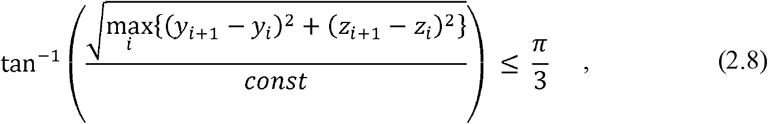

where

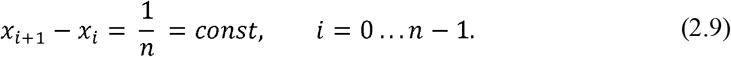

The additional constraint function (Equation 2.8) was implemented to ensure the smoothness of the optimised ligament paths and to prevent the ligaments from abruptly changing direction or intersecting the bone meshes between the path points. Hence the angular offset of any two subsequent ligament points could not exceed 60° relative to the ligament long-axis.

As with the ROM simulations, the above optimisation problem was solved using SQP [79]; see Supplementary Information.

### 2.7 Mobility simulation comparison

Resultant ligament lengths for each joint pose of the XROMM trials and ROM simulations were exported and subsequently analysed in Matlab 2023b. Outliers in estimated ligament lengths (i.e., instances in which the SQP optimiser failed to converge) were automatically detected and interpolated from the surrounding sine-corrected rotational data [16] using *scatteredInterpolant*. The resulting ligament lengths were compared to scaled dissection data and to the maximally sustainable strain observed in previous experimental data [59] and compared to maximal length estimates from the XROMM data. Joint poses in the ROM simulations for which any of the six simulated ligaments exceeded the strain tolerances were deemed impossible and, therefore, discarded from the osteological ROM dataset, as the ligaments would either have prevented the bones from attaining such a pose or would have torn. The resulting *in silico* soft tissue-constrained mobility estimates were then compared to the *ex vivo* mobility as determined through XROMM.

## 3. Results

### 3.1 Ex vivo *joint mobility*

Overall, 11 of 22 recorded trials were useable and fully analysed (RLP2, 2 trials; RLP3, 7 trials; RLP4, 2 trials; Figure S4; Table 1), approximating a total of four minutes of high-resolution motion data, resulting in 60,027 individual positions and orientations per bone. Eleven trials (50% of the collected data) had to be discarded due to suspected damage of the shoulder joint capsule; for both RLP2 and RLP4 potential damage occurred after two trials (Figure S4). The subsequent trials for these specimens were thus discarded from the shoulder dataset and any subsequent analyses (see [68]).

#### 3.1.1 Rotational movement

The three specimens showed similar constraints on joint mobility from their cadaveric manipulation, with their mobility envelopes mostly overlapping (Figure 2). Differences in overall excursion were relatively minor, with the specimens agreeing on individual mobility boundaries. Only two trials were collected prior to joint capsule damage for RLP2 and RLP4, thus their overall ROM observed (132,855.6°^3^ and 70,059.6°^3^ for RLP2 and RLP4, respectively) was substantially smaller than that of RLP3 (302,451.7°^3^). Due to the bulk of their individual ROM extent lying within the range seen in RLP3, these two specimens only marginally extended the combined XROMM volume to 488,189.2°^3^. The sine-corrected flexion/extension ROM of the combined taxa was almost exclusively constrained to positive values (i.e., movement with a laterally oriented humerus, as negative values indicate a medially directed humerus; Figure 2).

**Figure 2.**
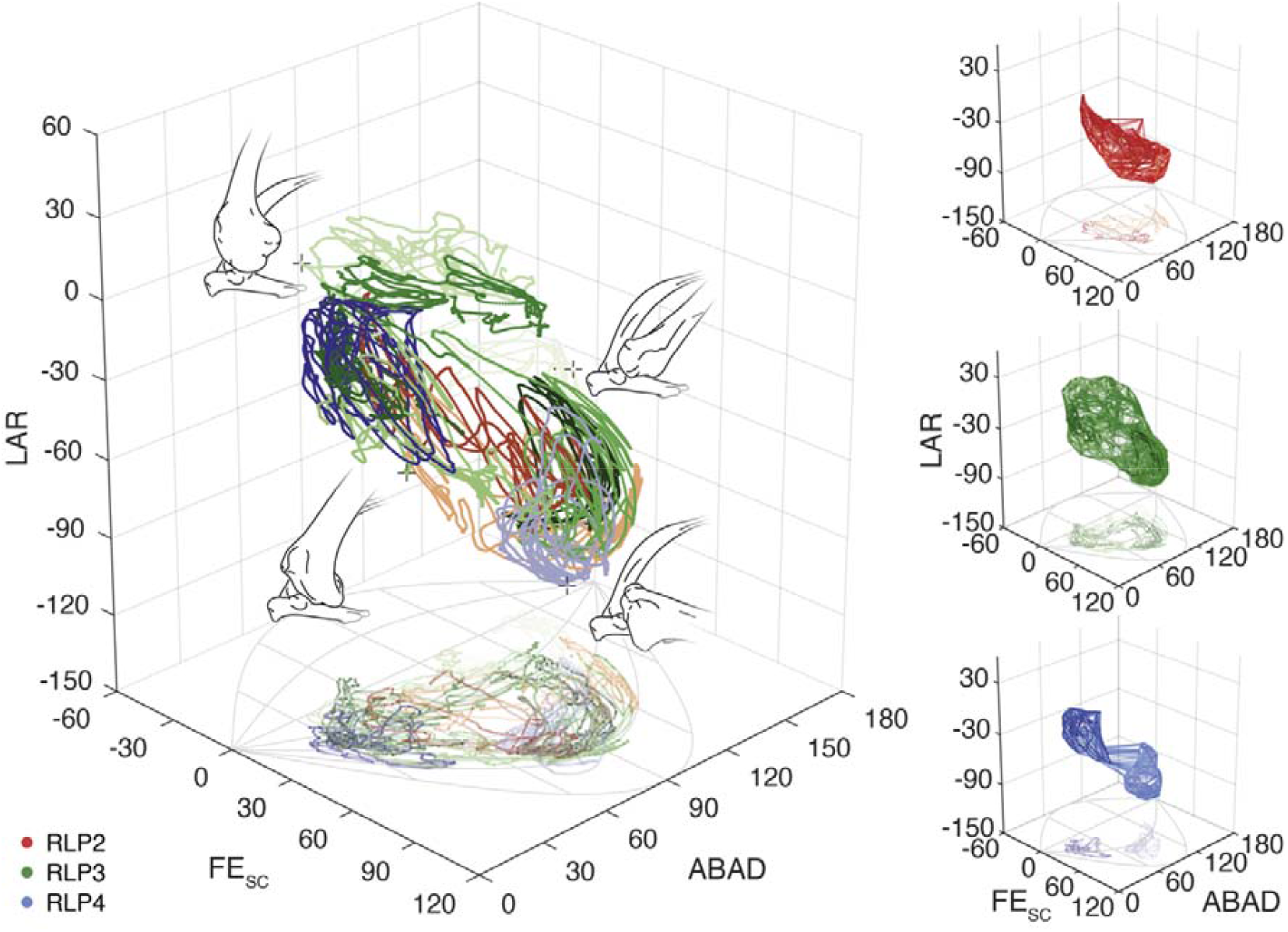
*Ex vivo* rotational mobility of the different RLP specimens. Combined and individual rotational joint mobility of the different specimens in the sine-corrected (SC) space of the *Two-Neutral Spherical Rotation System* [16]. Illustrations show a selection of representative joint orientations of the humerus relative to the scapulo-coracoid with their corresponding data points denoted by crosses. Individual trials are colour-coded from dark to light for each specimen. Abbreviations: FE, flexion/extension; ABAD, abduction/adduction; LAR, long-axis rotation.

#### 3.1.2 Translational movement

Cadaveric manipulation revealed substantial sliding (i.e., translational movement) in the RLP shoulder joint (Figure 3; Figure S5). Due to the large magnitudes of joint translations, the superimpositions of the ACSs (as determined by the fitted shapes to the glenoid and humeral head articular surfaces) were a poor approximation of the functional joint centre (Figure S6), unlike what has been suggested for other joints, such as the hip joint of other archosaurs [12] or the hip joint in hominins [56].

**Figure 3.**
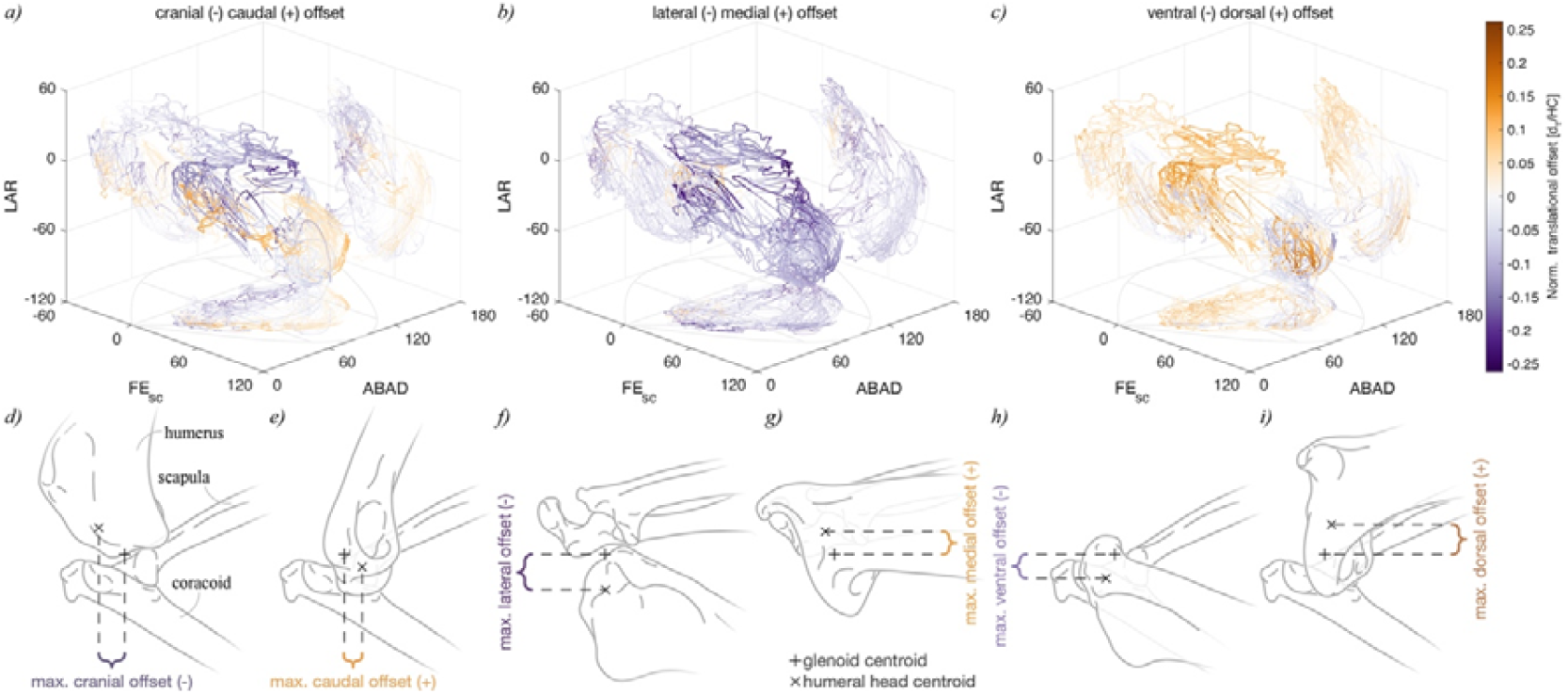
*Ex vivo* translational joint mobility in the RLP shoulder joint. Size-normalised craniocaudal (a,d,e), mediolateral (b,f,g) and dorsoventral (c,h,i) offsets between the humeral head and glenoid centroids. Individual rotational poses are colour-coded by the corresponding values of respective translational DOFs. Illustrations (d-i) demonstrate shoulder poses with the maximal translational offsets in lateral (d,e,h,i) and dorsal (f,g) views. RLP shoulder mobility indicates only limited coupling between rotations and translations across their range of motion, as large translational deviations were not restricted to specific rotational areas and *vice versa*. Abbreviations: d_T_, translational displacement; HC, humeral circumference.

Substantial interactions within and between translational and rotational DOFs were observed, with certain joint positions displaying exhibiting higher variability in joint orientations than others. Large translational excursions were not limited to specific orientations of the humerus relative to the glenoid (Figure 3). Rotational and/or translational minima or maxima of individual DOFs were never attained simultaneously, similar to previous studies [33,38,39]. There was no apparent coupling between translational and rotational DOFs in the shoulder joint of the RLP specimens (i.e., extremes in rotations are not constrained to extremes in translations and *vice versa*; Figure 3) and translational extremes were distributed evenly across the rotations, unlike reported for other tetrapods [14,37].

#### 3.1.3 Ligament length estimates

The calculated ligament lengths from the XROMM data approximated the scaled dissection data (Figure 4; Supplementary Table S1) for all specimens, while highlighting differences in observed strain ranges among individual ligaments. Ligament length estimates stayed within or below the range of maximally feasible strain in experimental data (17%–47%) of the acrocoracohumeral ligament (LAcH) [59]. The coracohumeral ligament (LCoHd) exhibited the most strain in the cadaveric XROMM data, exceeding the mean failure strain of the LAcH (26% [59]) in both RLP2 and RLP3, with RLP4 remaining just below the threshold. The lateral (LScHl) and posterior (LScHp) scapulohumeral ligaments never reached a taut state during the *ex vivo* experiments in two specimens (RLP3 and RLP4 for the former, and RLP2 and RLP4 for the latter; Figure 4). The fact that RLP2 and RLP4 never reached a taut state for the LScHp might be due to those specimens’ generally smaller ROM extent (Figure 2) which did not fully cover the total ROM in the shoulder before damage to the joint capsule occurred and, therefore, did not stretch the ligaments to their full extent. The greatest strain range was observed in the intracapsular ligament (LIcCa), with the distance between origin and insertion ranging from 5.4% of resting length (−94.6% strain) to 139.5% (39.5% strain).

**Figure 4.**
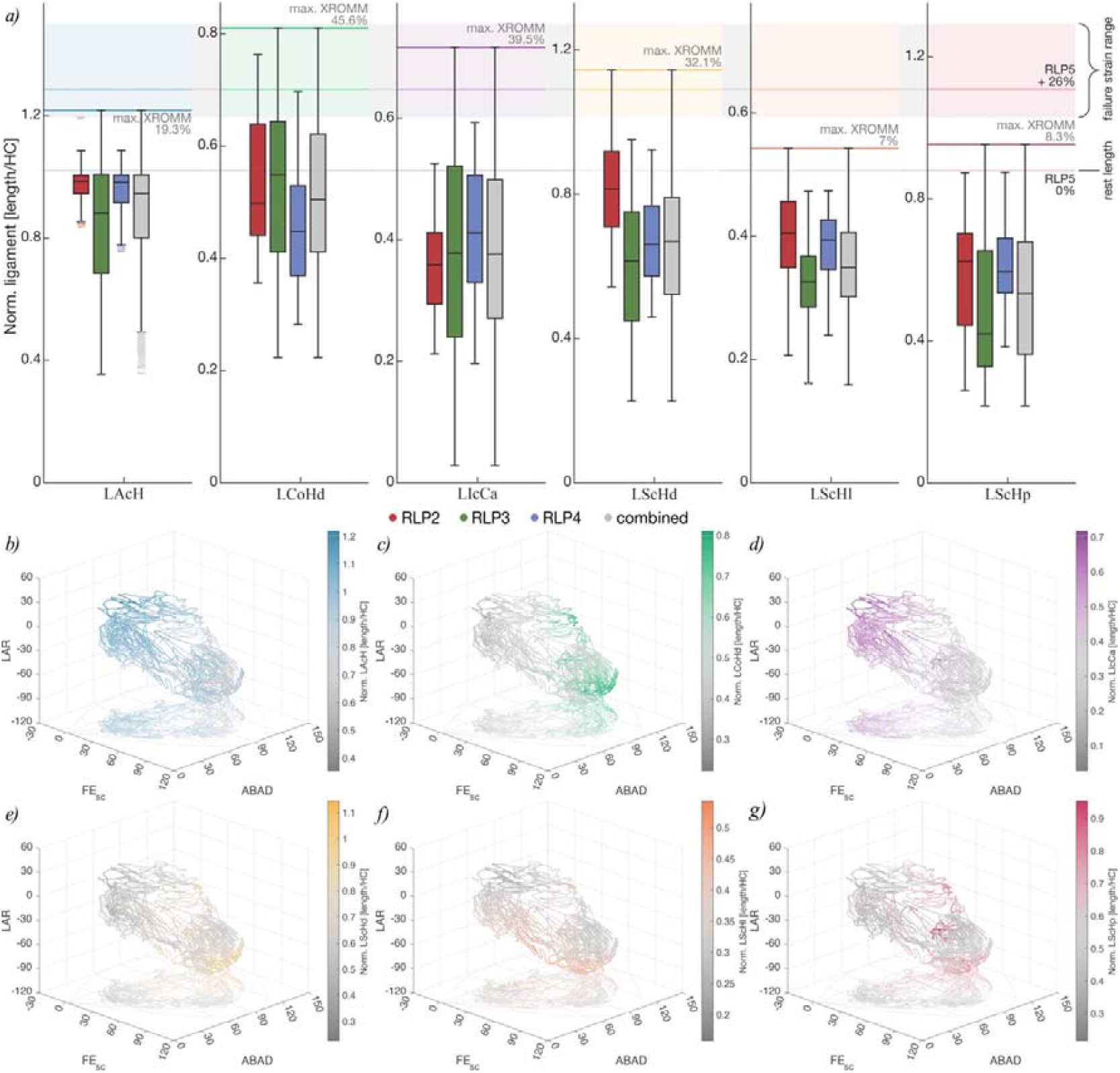
Normalised ligament length estimates based on XROMM trials. (a) Comparison between size-normalised ligament lengths from XROMM data and scaled dissection data with experimental failure strain ranges of the acrocoracohumeral ligament [59]. (b-g) Normalised ligament lengths calculated from the XROMM data in relation to joint excursion in the sine-corrected *Two-Neutral Spherical Rotation System*. Colours correspond to ligaments in (a).

The greatest ligament strains were observed in abducted (elevated, ABAD <90°) joint poses with positive axial rotation (LAR) for the LAcH and LIcCa, whereas the opposite was true for the LCoHd and the three LScH (adducted/depressed ABAD >90°; Figure 4b-g). Note that the *Two-Neutral Spherical* transformation does not directly correlate with motion around the cardinal axes due to embedded axial rotations and is thus less intuitive; however, it is able to prevent issues related to gimbal lock and thus preserves relative distances between individual joint poses in 3D [16].

### 3.2 In silico *joint mobility*

#### 3.2.1 Osteological ROM

The osteological ROM overestimated the *ex vivo* joint mobility captured via XROMM by an order of magnitude across all specimens (Table 2). The pose space generated by the ROM simulations initially resulted in many osteologically feasible, yet physiologically unlikely joint poses if soft tissue constraints were considered (e.g., ligaments).

**Table 2.** Simulation results and comparison with XROMM data.

| Specimen | Dataset | Volume [<sup>o</sup>3] | Missed [<sup>o</sup>3] | Missed [%] |
| --- | --- | --- | --- | --- |
| RLP2 | ROM | 3,550,553.25 | 30,144.40 | 5.59 |
|  | Dissection | 1,573,886.51 | 162,207.39 | 30.08 |
|  | XROMM | 1,919,651.02 | 92,909.87 | 17.23 |
|  | Full | 6,100,591.27 | 26,407.15 | 4.90 |
| RLP3 | ROM | 3,202,769.72 | 10,413.26 | 1.93 |
|  | Dissection | 2,841,991.28 | 96,699.85 | 17.93 |
|  | XROMM | 3,081,498.41 | 9,337.84 | 1.73 |
|  | Full | 5,993,494.98 | 2,944.38 | 0.54 |
| RLP4 | ROM | 3,051,695.43 | 585.96 | 0.11 |
|  | Dissection | 2,048,592.97 | 129,366.39 | 23.99 |
|  | XROMM | 2,212,443.72 | 79,421.31 | 14.73 |
|  | Full | 6,156,200.81 | 301.59 | 0.06 |
| All | XROMM | 539,327.29 | - | - |

The osteological ROM simulations of all specimens failed to capture portions of the experimental movement envelopes, especially joint poses with a combination of negative FE_SC_ and LAR angles (Figure 5d), missing from 585.96°^3^ (RLP4) to 26,407.15°^3^ (RLP2), respectively representing 0.06% and 4.90% of the total range of cadaveric joint mobility. While the full range was not recovered in these osteological simulations, the missing areas were predominantly located at the border of cadaveric joint mobility and may represent an artefact of the sampling resolution, especially in RLP3 and RLP4, where the missing areas remain below 1%. The ROM simulations were sampled at 5-degree intervals [31,35] due to a trade-off between accuracy and processing time, as joint poses and the resulting processing time both scale exponentially [14,35] with a decreasing sample interval (on average 0.613 seconds per joint pose for a random subset of 10,000 joint orientations on an Apple M2 Pro @3480 MHz on a single core). Given the small percentages of missed data in those two specimens (<1%), any missed joint poses would almost certainly have been recovered with a higher sampling resolution and are thus deemed negligible. Decreasing the sample interval (and thus exponentially increasing the number of joint poses) would have come at the expense of impractical computational runtimes for these simulations.

**Figure 5.**
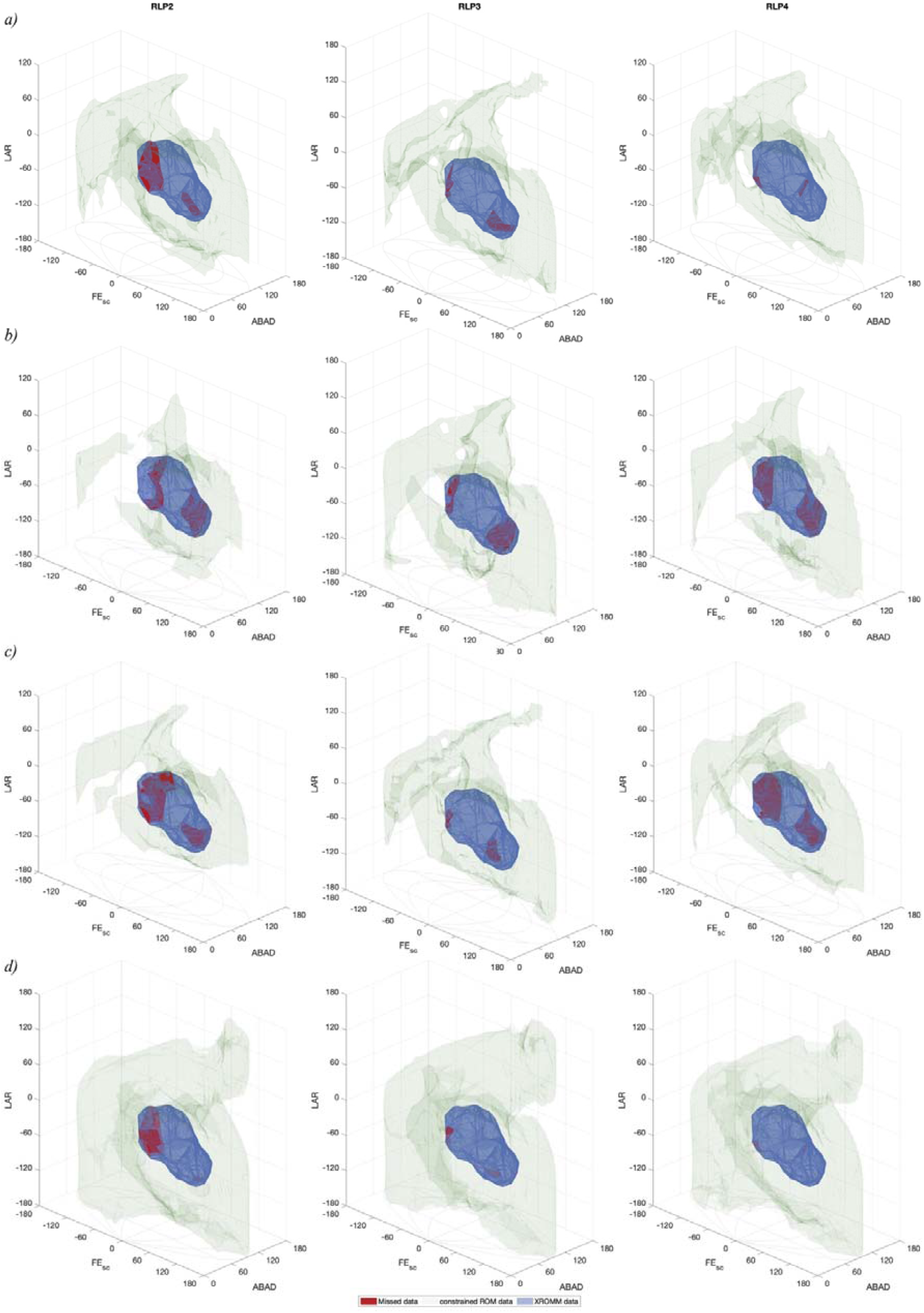
Comparison between cadaveric joint mobility, osteological and ligament-constrained ROM simulation data. (a) ROM-constrained simulation, in which the ligament thresholds were based on the interpolated maximal lengths for each ligament based on the ROM simulation dataset across all XROMM poses. (b) Dissection data-constrained ROM simulations using scaled and ‘strained’ dissection data of RLP5 (i.e., +26% strain; see Table 2, Supplementary Table S1). (c) XROMM data-constrained ROM simulations. (d) Full osteological ROM without any soft tissue constraints. Red volumes represent missed poses, green volumes are the constrained ROM simulations and blue volumes are the combined *ex vivo* joint mobility.

#### 3.2.2 Ligamentous constraints on mobility

The constraints imposed by the simulated ligaments substantially reduced osteological joint mobility estimates (Figure 5). While the scaled and ‘strained’ dissection data yielded the least mobility overall, it was a poor approximation of the *ex vivo* mobility due to missing substantial portions of cadaveric movement (missing 17.93–30.08% of total *ex vivo* joint mobility; Table 2).

Ligament strain thresholds used to constrain the ROM volume are in Supplementary Table S1. The ‘ROM’ dataset represents the constrained dataset based on interpolated maximal lengths across all XROMM poses for each ligament in the ROM simulation dataset; ‘XROMM’ represents the constrained dataset based on maximal ligament lengths directly calculated from the XROMM dataset; ‘Dissection’ represents the constrained dataset based on scaled dissection values with an additional strain factor of 26% (mean failure strain from experimental data of the acrocoracohumeral ligament [59]); the ‘Full’ dataset represents the unconstrained ROM dataset. Note that differences between the ROM and XROMM datasets are due to differing translational positions for identical rotational poses which directly influence the calculated length of any given ligament. Maximal ligament lengths calculated from the ROM datasets tend to be greater than those from the XROMM datasets (see Supplementary Table S1).

Differences in the missed volumes between the full osteological ROM and the ROM-constrained estimates (i.e., the maximal interpolated values from the ROM ligament dataset across all XROMM coordinates) were relatively minor (<1.5%), only marginally increasing the number of missed poses (Table 2), whereas the soft tissue constraints substantially reduced overall joint mobility by 47.1% (RLP2), 39.3% (RLP3) and 32.6% (RLP4). Due to differences in joint translations between the XROMM poses and the optimised ROM positions, the calculated maximal ligament lengths for the latter often exceeded the former (except for LScHp in RLP2; see Supplementary Table S1). Therefore, the XROMM-constrained ROM estimates often missed substantially more cadaveric joint mobility than the ROM-constrained estimates, apart from the case of RLP3, for which the XROMM-constrained mobility estimates missed less of the *ex vivo* poses than the ROM-constrained estimates.

## 4. Discussion

### 4.1 Importance of the absence of a functional joint centre

The centroids of either shape fitted to the articular surfaces failed to predict the shoulder joint centre position accurately, even though the mean joint centre was mostly positioned within these shapes (Figure S5). This has fundamental implications for reconstructing joint mobility of extinct taxa. Considering that extensive translational movement is evident in the bird shoulder [59,60,78] and present in other animal joints [9,37], the functional (mean) joint centre position cannot be reliably inferred from the preserved bony articular surfaces via shape-fitting. Primitive shape fitting is a useful tool for standardising joint and bone orientations [53,73]; however, the resulting joint axes and joint centre position may not necessarily represent the true functional joint axes or their centroids for complex joints, but rather a standardised reference frame enabling comparisons among multiple specimens. Translational DOFs must, therefore, be included in any such comparison of joint mobility estimates [33], otherwise the exclusion of possible but missed joint poses may result in erroneous conclusions and/or functional inferences. In a joint as open as the avian shoulder joint [60,78], translations need to be considered to obtain accurate mobility estimates [33,36,56,59]. Therefore, joint mobility simulations using only a single joint centre (or multiple rigid ones) are inadequate for accurate estimates of joint mobility. Only truly dynamic joint centres, constrained via consistent articular congruence as presented here (or similar approaches presented elsewhere; e.g., see [14,16,83]), enable accurate osteological mobility estimates.

### 4.2 Soft tissue influences on avian shoulder joint function

The avian shoulder joint is subject to large forces during flight which are mitigated by a suite of ligaments that prevent dislocation of the joint during movement [60,78,84]. Whereas the LAcH is the main ligament preventing ventral dislocation of the shoulder, counteracting and transmitting the pulling force of the pectoralis muscle through the strut-like coracoids [84], it also secondarily restricts excessive abduction of the humerus in combination with the LIcCa. Excessive adduction of the humerus is prevented by the LCoHd and the LScHd, and those ligaments additionally support the suspension of the humerus during the down stroke, whereas the LScHl and LScHp prevent excessive extension/protraction. As with most collateral ligaments, the avian shoulder ligaments do not only prevent excessive rotational movements but also translations beyond natural joint mobility and ensure that the joint surfaces stay in articulation during the considerable sliding movements present in the avian shoulder joint. The especially osteologically unconstrained and open hemi-sellar morphology of the avian glenoid [60,78] differs from the more constrained shoulder joints of other (quadrupedal) tetrapods [14,34,50], where joint stability is more desirable and constraints are necessitated to counteract and support the animal’s body weight. In our experiments, the hemi-sellar glenoid’s articular cartilage constrained translational movement mostly to an axis perpendicular to it (Figures 3,S5), as was hypothesised by Sy [78], increasing rotational movement of the humerus during flapping motion, whereas the ligaments facilitate constant contact within the shoulder capsule.

Individual ligaments appear to show differences in their elasticity (Figure 4), with the LAcH being relatively stiffer than the other five, with the observed strain remaining below 20% in all specimens, whereas the LScHd, LIcCa and LCoHd all exceeded 30% strain during cadaveric movement. This could indicate potential differences in the composition and mineral content of these ligaments (e.g., see [85]) and that some may be able to passively store and release elastic energy for usage during the wing stroke, which deserves future investigation.

Furthermore, given that ligaments are an integral component of the force balance within the avian shoulder joint during flight [59], simulating ligamentous forces and calculating their strain could be an important factor in improving musculoskeletal models of avian wing and shoulder function (e.g., [66,86,87]) that has previously been overlooked.

## 5. Conclusions

This study represents a significant advance in our ability to implement and infer soft tissue constraints on joint mobility estimates in both living and extinct taxa to overcome some previous challenges set by translational joint sliding. The implementation of the algorithms initially outlined by Marai and colleagues [75,76] into Autodesk Maya directly integrates them into the most commonly used software package for joint mobility estimation and skeletal animation in non-human vertebrates [10,12,15,19,25,26,28–31–34,38,40,42,43], and presents a significant improvement and expansion of the existing pipeline for investigating form-function relationships of vertebrate joints. Joint ROM simulations are no longer limited to physiologically implausible static joint centres but approximate the true behaviour of joints and their articulation using contact-based optimisation. Such simulations intrinsically describe 6 DOFs in their motion. By simulating soft tissues, osteological ROM can be constrained to a more functionally informative mobility envelope, thus enabling more accurate inferences through the exclusion of physiologically infeasible joint poses.

Understanding the function of the avian shoulder joint is fundamental to understanding the avian wing stroke. Evidently, the bird shoulder cannot be approximated by an idealized ball and socket joint. Indeed, substantial sliding and/or rolling occurs within the hemi-sellar avian shoulder joint in order to achieve certain joint poses (Figures 2,3) [59,60,78]. Thus, hemi-sellar joints are not necessarily less mobile than ball-and-socket joints, but rather employ translations to increase mobility. The reorientation of the glenoid to face dorsolaterally in birds, rather than posterolaterally as in non-avian reptiles, accommodates the avian wing stroke [60]. The importance of this evolutionary reorientation is reinforced by our findings for the mobility of the partridge shoulder presented here. Reorientation of this joint directly facilitated the large degrees of abduction/elevation evident in our cadaveric joint mobility results (Figure 2), such that the wing can be raised far above the shoulder. This ability may represent a fundamental adaptation facilitating takeoff from the ground and the active, flapping flight of many extant birds. Such movement is not permitted with a laterally facing glenoid like that of extant non-avian reptiles [37], and non-avian dinosaurs [60].

Our approach will help facilitate justified functional and evolutionary inferences of joint mobility in taxa that are extinct or otherwise impossible to study with *in vivo* data, adding to an increasing methodological repertoire to help researchers decipher articular form–function relationships and their evolutionary history. Simulated soft tissues constrain the overexpansion of osteological mobility estimates caused by sliding within joints through the exclusion of physiologically unfeasible joint poses and approach cadaveric mobility more closely. Our study’s methodological innovations will allow functional interpretations of joint mobility data and improve palaeobiological inferences regarding joint form and function throughout the evolutionary history of vertebrates.

## Author contributions

OED, JRH and DJF conceived and designed this study. OED and SEW collected the experimental data. OED and VLB wrote and implemented the Python code. OED segmented the CT data, tracked the trials, analysed and interpreted the data, prepared the figures, and wrote the manuscript. All authors contributed to editing and revisions of the manuscript and approved the final version.

## Acknowledgements

We thank Andrew Cuff, Peter Bishop and Armita Manafzadeh for helpful discussions regarding the setup of the biplanar X-ray. We further thank Keturah Smithson for conducting the μCT scans of the partridge specimens at the University of Cambridge Tomography Centre. We thank Pasha van Bijlert for fruitful discussions regarding the implementation and setup of the cost functions. We are grateful to Emily Mitchell and Sophie Regnault for comments and discussions that improved the manuscript.

## Data accessibility

The datasets generated and/or analysed during the current study are included in this published article (and its Supplementary Information files). The Python and Matlab code can be found in the following Zenodo and Github repositories: https://doi.org/10.5281/zenodo.15442127 and https://github.com/OliverDemuth/MayaSignedDistanceFields. The μCT scans of RLP2-4 and their XROMM trial data are available from the following Zenodo repositories: https://doi.org/10.5281/zenodo.15436559 (CT data), https://doi.org/10.5281/zenodo.15373685 (exemplary XROMM trial) and https://doi.org/10.5281/zenodo.15437558 (marker trajectories and ROM datasets).

## Declaration of AI use

We have not used any AI-assisted technologies in creating this article.

## Ethics

This work did not require ethical approval from an animal welfare committee.

