## Supplementary_Information for "Soft tissue constraints on joint mobility in the avian shoulder"

**Supplementary Figures**

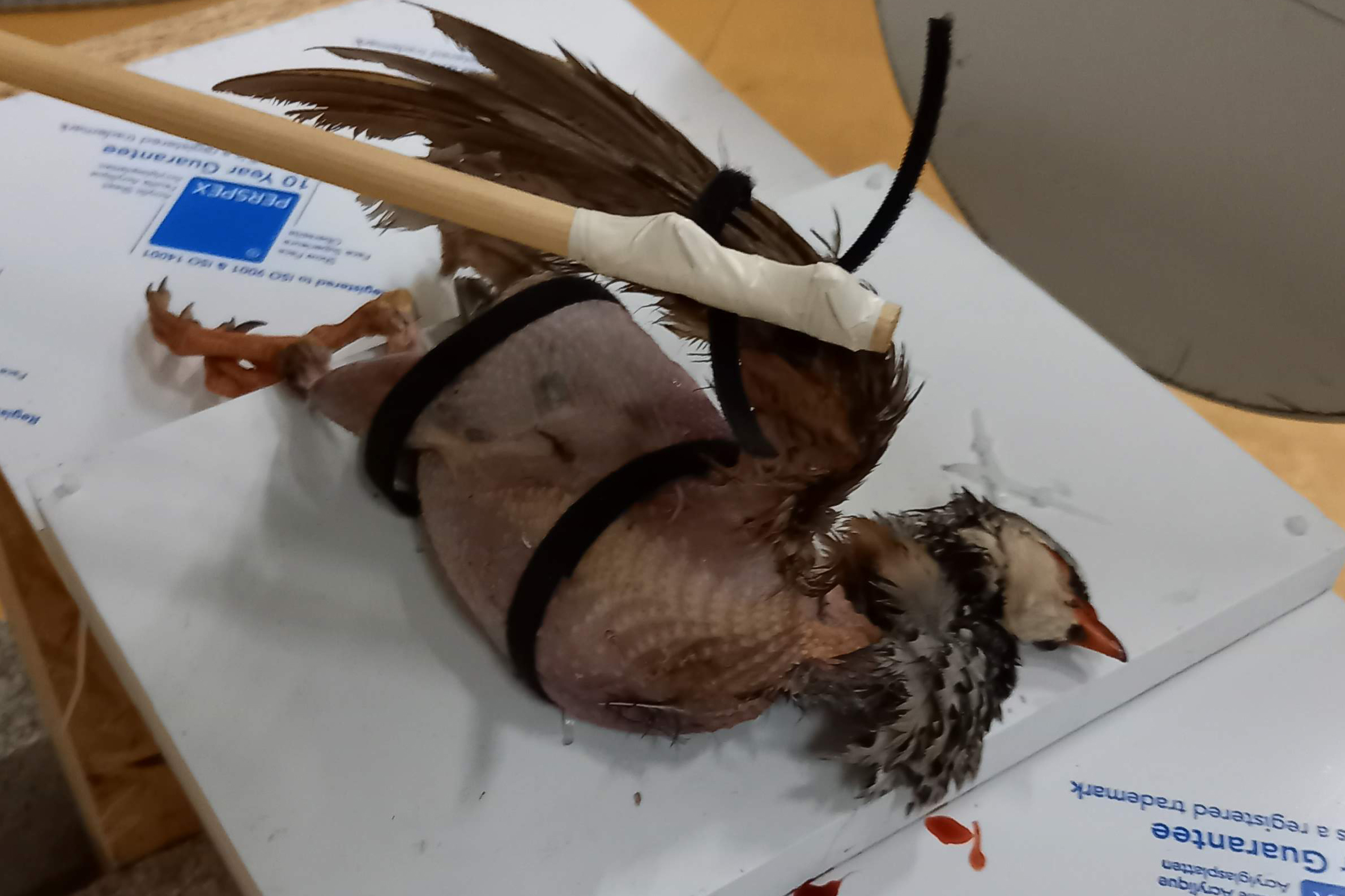

**Figure S1. Velcro harness and setup for the XROMM experiments.** Specimen RLP2 rests on the foam board within the capture area of the biplanar fluoroscopes. It is secured with a Velcro harness around the body and on the wing, which was loosely fitted to minimise potential damage to the bird’s soft tissues due to the large lever arm of the manipulator rod.

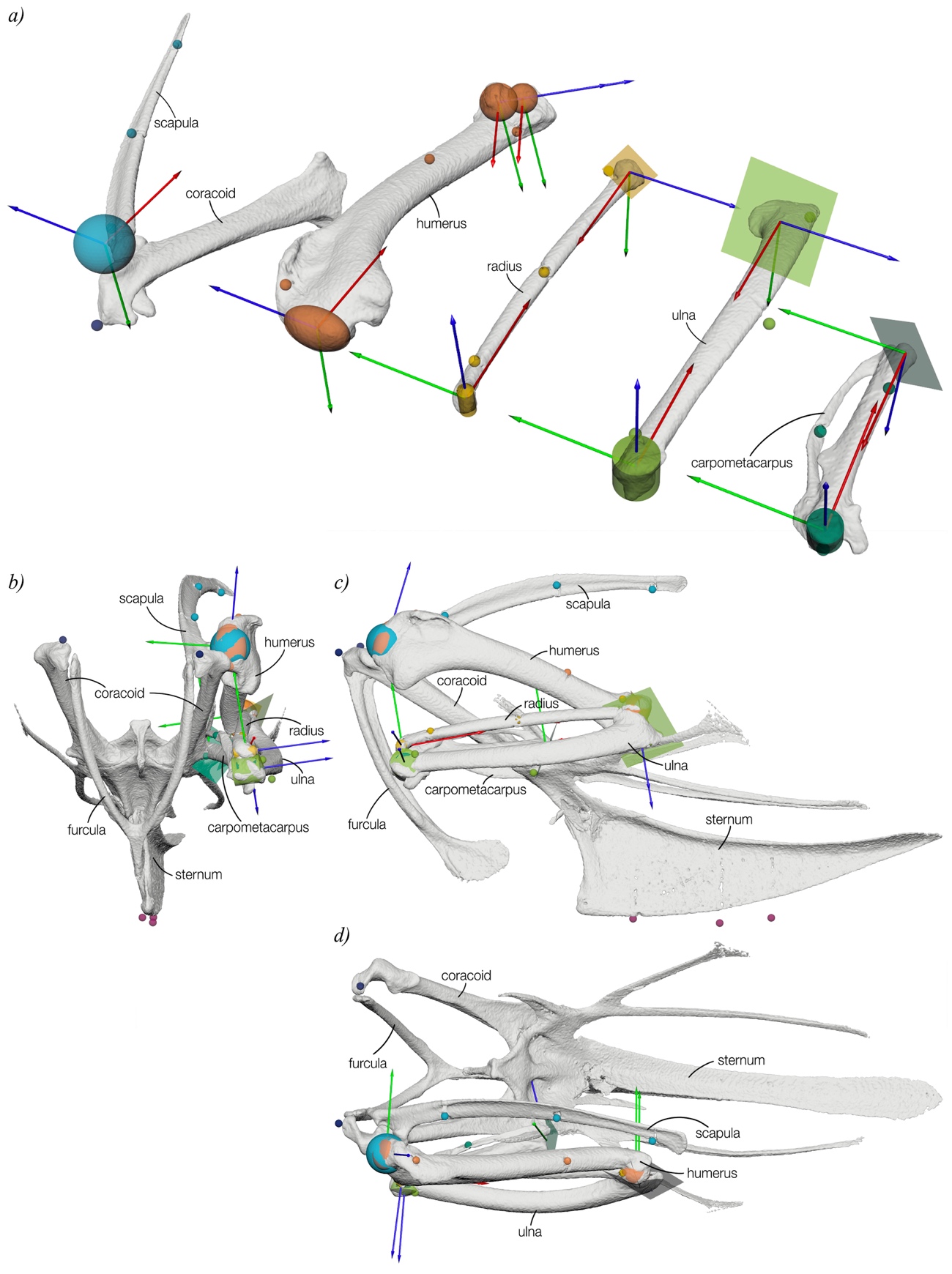

**Figure S2. ACS and JCS setup for RLP specimens.** (a) Joint axes definitions based on fitted shapes. Exemplar reference pose with collapsed wing in cranial (b), lateral (c) and dorsal (d) view of RLP3 in which all joint angles were set to 0°.

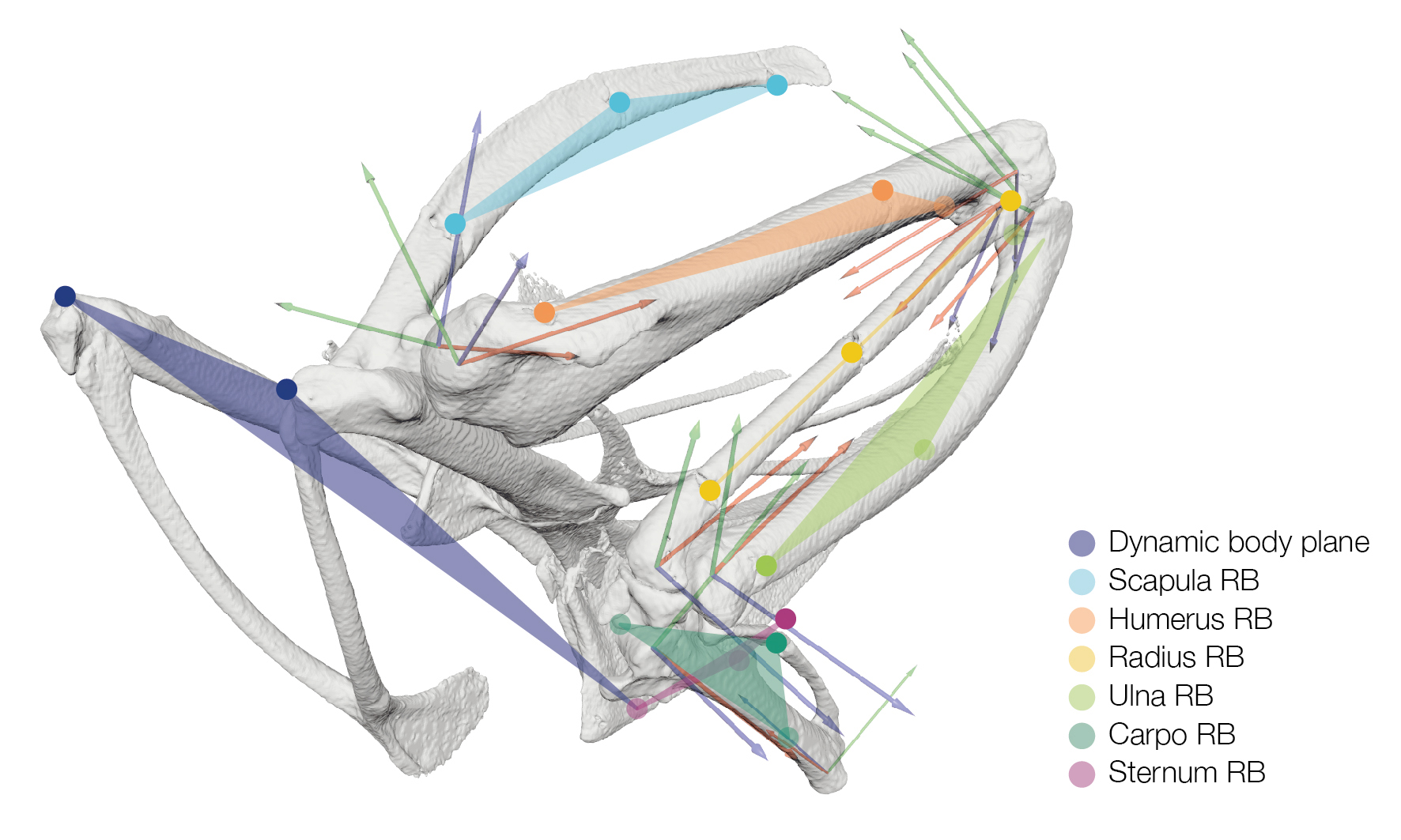

**Figure S3. Rigid body definition for RLP specimens.** Rigid bodies (RB) for each bone were defined by three points. ACSs and JCSs were then animated based on bead movement from the XROMM trials.

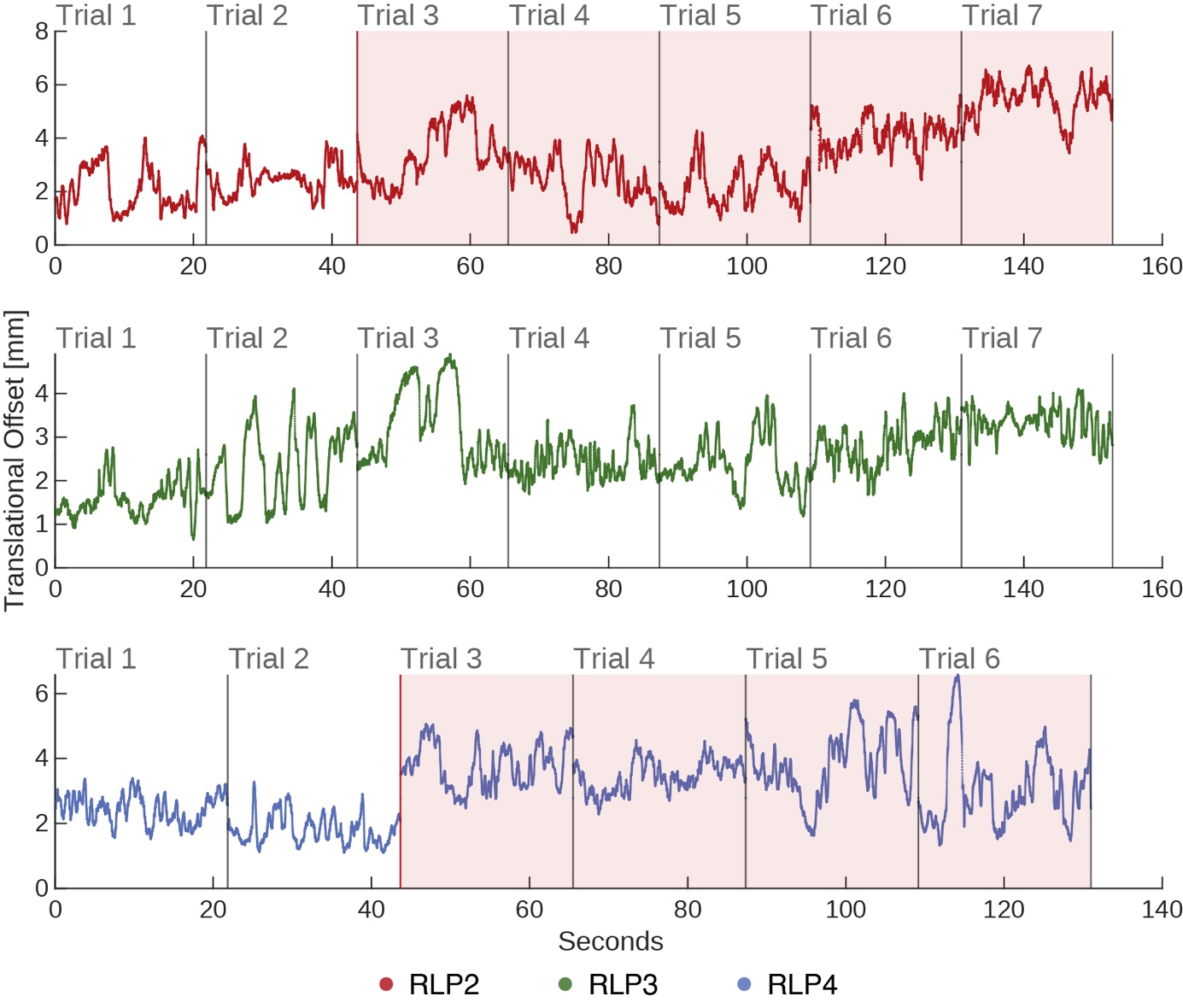

**Figure S4. Offset of the humeral head centroid relative to the glenoid centroid across all trials.** Red lines in the traces for RLP2 and RLP4 indicate the point at which damage to the shoulder joint capsule occurred. All subsequent trials for these specimens were discarded. Trials 7 and 8 of RLP4 were not processed as the four previous trials already indicated substantial pathological sliding.

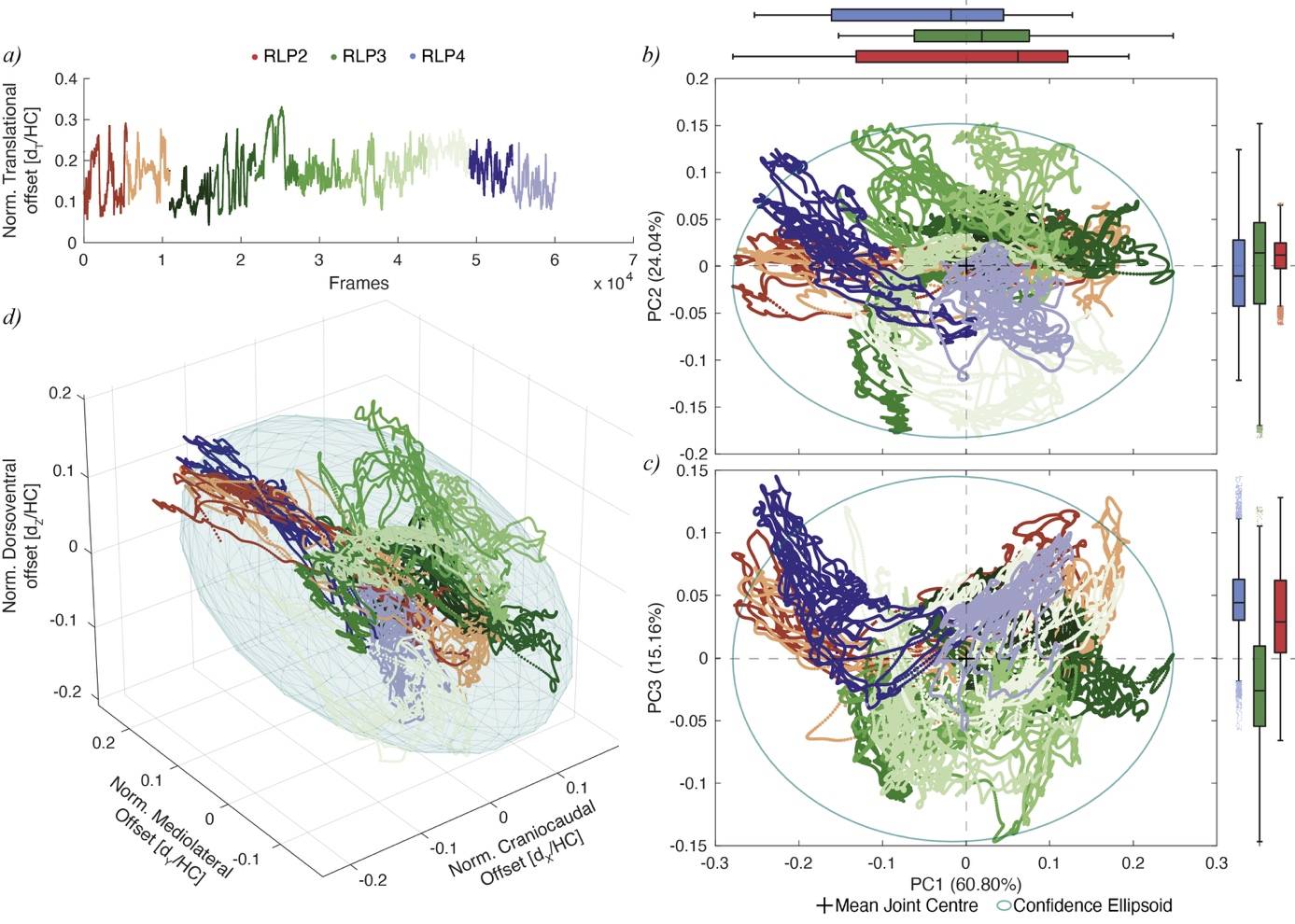

**Figure S5. *Ex vivo* translational joint mobility in the RLP shoulder joint.** (a) Size-normalised distance between the proximal humeral ACS and the glenoid mean joint centre position for all frames. Individual trials are colour-coded from dark to light for each specimen with marginal box plots describing their translational range along the PC axes. (b,c) PCA across all joint poses with inferred confidence ellipsoid and the calculated mean joint centre position. (d) Point cloud of the joint position and three-dimensional representation of the confidence ellipsoid in the coordinate system of the glenoid joint.

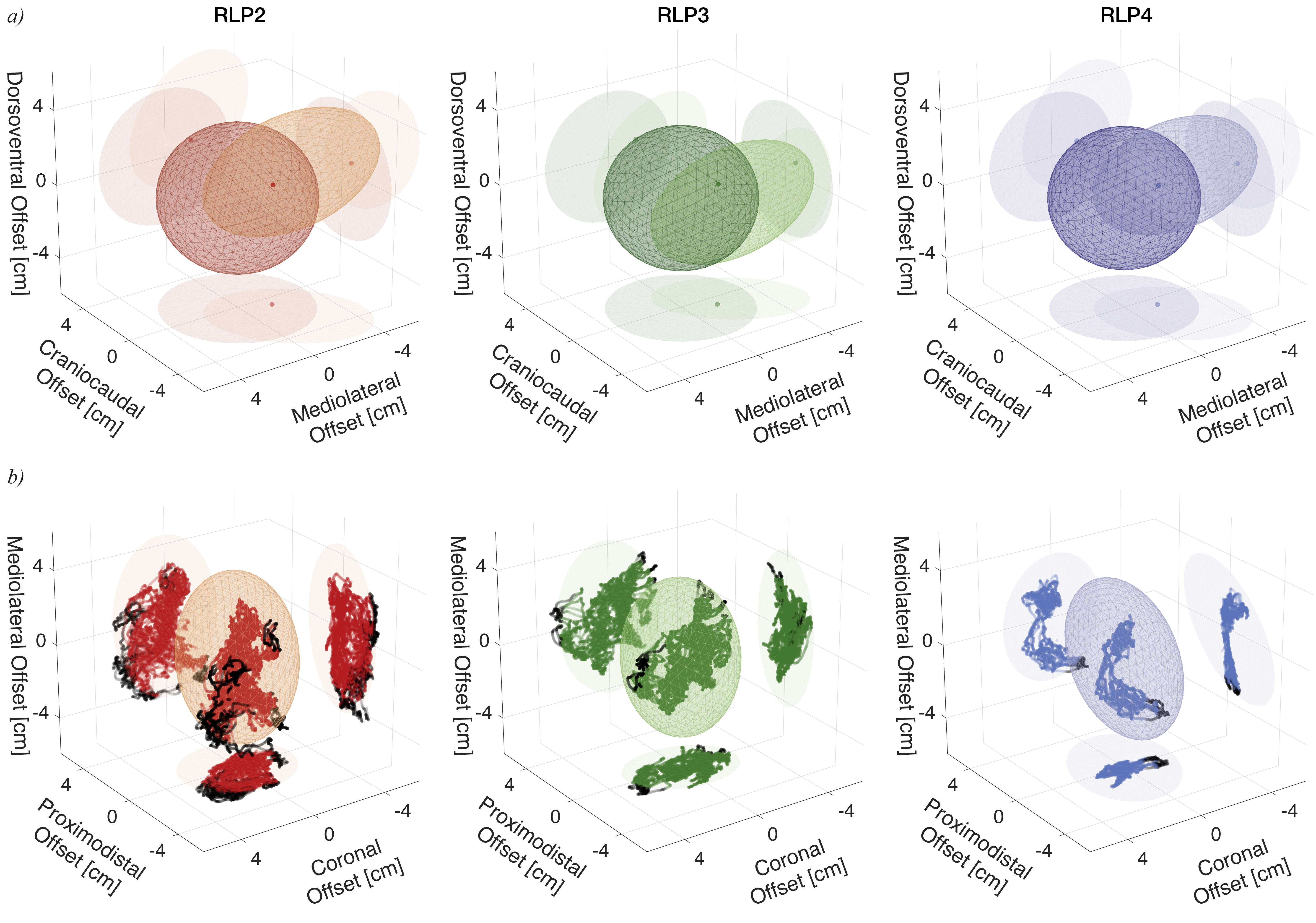

**Figure S6.** **Mean joint centre position relative to fitted primitive shapes in the RLP shoulder joints.** (a) Position of the individual mean joint centre and ellipsoid fitted to the humeral head (in the CT scan position) relative to the glenoid ACS and its fitted sphere. The mean joint centre lays within the intersecting volume of the glenoid sphere and the humeral ellipsoid in all three specimens. (b) Relative positions of the mean joint centre to the humeral ACSs. Note, while the mean joint centre remains stationary within the sphere (a), it dynamically shifts in relation to the humerus coordinate system and sometimes leaves the humeral head ellipsoid (b), as indicated as black dots. Shadows of the shapes and points are projected onto the individual planes to better illustrate their positions in 3D.

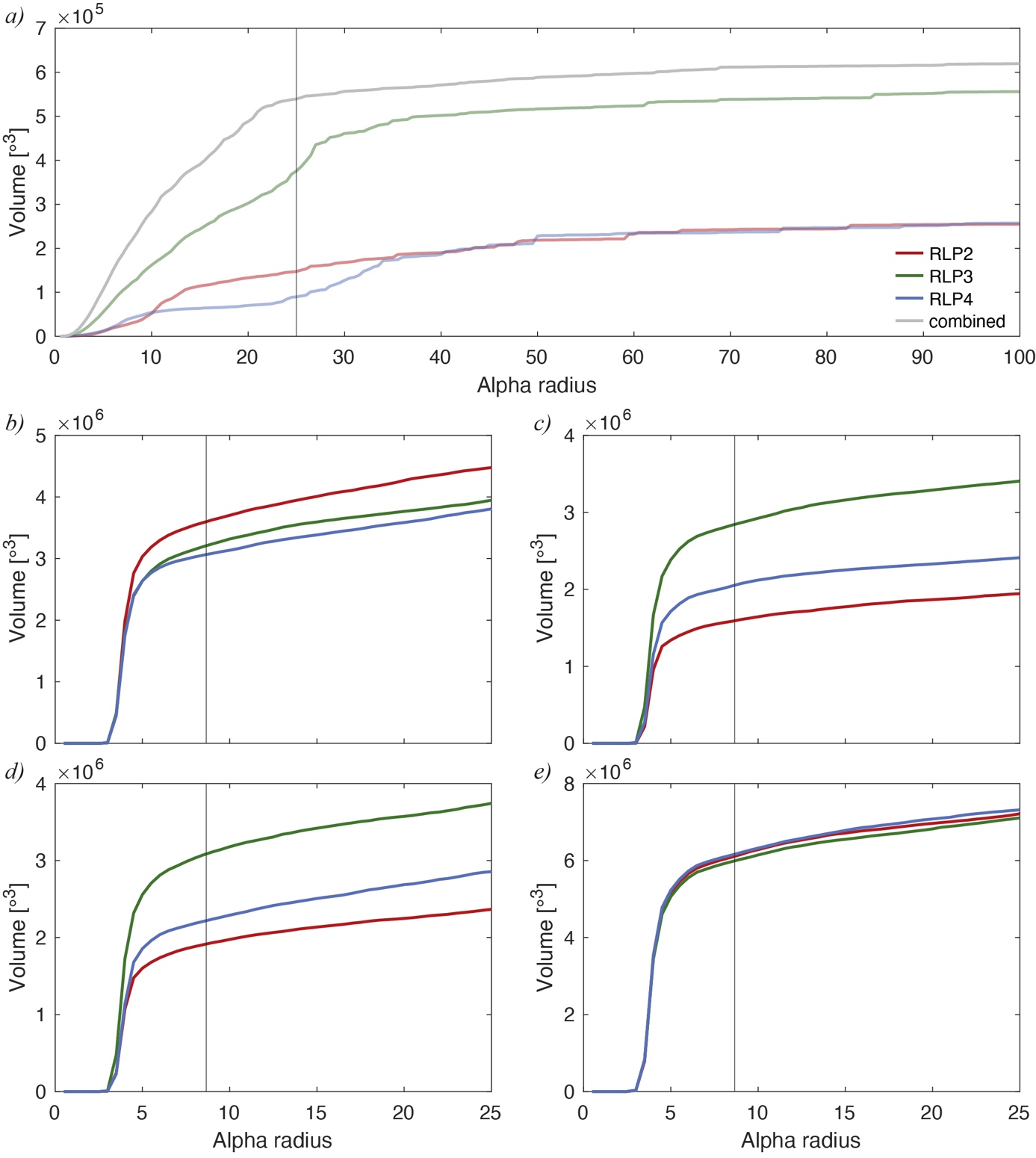

**Figure S7.** **Sensitivity analysis of alpha shape radii.** Volumes of *ex vivo* (a) and *in silico* (b-e) joint mobility analyses. (a) Cadaveric mobility volumes for individual specimens and combined XROMM data. (b) ROM-constrained simulation, in which the ligament thresholds were based on the interpolated maximal lengths for each ligament based on the ROM simulation dataset across all XROMM poses. (c) Dissection data-constrained ROM simulations using scaled and ‘strained’ dissection data of RLP5 (i.e., + 26% strain; see Table 2, Supplementary Table S1). (d) XROMM data-constrained ROM simulations. (e) Full osteological ROM without any soft tissue constraints. Vertical lines indicate the optimal alpha (α) radii used for analyses, which determine how tightly the concave hull wraps around the point cloud. The radii were set at 25 (a) for the experimental *ex vivo* XROMM data and $5\sqrt{3}$ (b-e) for the *in silico* simulations (i.e., the diagonal of the cube with a side length of 5, which was the sampling interval of the simulations). The optimal α radii avoided the sharp decrease in volume caused by smaller values and preceded the gradual but substantial inflation of the volume.

**Supplementary Tables**

**Supplementary Table S1. Ligament length estimates and comparison.**

| Specimen | Body mass [g] | HC [mm] | Ligament | Scaled dissection data [mm] | Dissection data + strain [mm] | ROM [mm] | scaled XROMM [mm] |
| --- | --- | --- | --- | --- | --- | --- | --- |
| RLP2 | 527 | 14.06 | LAcH | 14.36 | 18.09 | 20.31 | 17.12 |
|  |  |  | LCoHd | 7.82 | 9.86 | 12.45 | 11.39 |
|  |  |  | LIcCa | 7.23 | 9.11 | 12.16 | 10.08 |
|  |  |  | LScHd | 12.18 | 15.35 | 22.24 | 16.09 |
|  |  |  | LScHl | 7.13 | 8.98 | 9.49 | 7.63 |
|  |  |  | LScHp | 12.38 | 15.59 | 13.38 | 13.40 |
| RLP3 | 492 | 14.83 | LAcH | 15.14 | 19.08 | 18.38 | 18.06 |
|  |  |  | LCoHd | 8.25 | 10.40 | 11.65 | 12.01 |
|  |  |  | LIcCa | 7.62 | 9.61 | 10.31 | 10.63 |
|  |  |  | LScHd | 12.85 | 16.19 | 18.74 | 16.97 |
|  |  |  | LScHl | 7.52 | 9.47 | 8.72 | 8.05 |
|  |  |  | LScHp | 13.05 | 16.45 | 14.58 | 14.14 |
| RLP4 | 420 | 13.49 | LAcH | 13.78 | 17.36 | 19.03 | 16.43 |
|  |  |  | LCoHd | 7.51 | 9.46 | 11.14 | 10.92 |
|  |  |  | LIcCa | 6.94 | 8.74 | 10.74 | 9.67 |
|  |  |  | LScHd | 11.69 | 14.72 | 20.38 | 15.44 |
|  |  |  | LScHl | 6.84 | 8.62 | 7.40 | 7.32 |
|  |  |  | LScHp | 11.88 | 14.96 | 14.20 | 12.86 |
| RLP5 | 421 | 14.2 | LAcH | 14.5 | - | - | - |
|  |  |  | LCoHd | 7.9 | - | - | - |
|  |  |  | LIcCa | 7.3 | - | - | - |
|  |  |  | LScHd | 12.3 | - | - | - |
|  |  |  | LScHl | 7.2 | - | - | - |
|  |  |  | LScHp | 12.5 | - | - | - |

Ligament length estimates were scaled based on humeral circumference (HC). Strained dissection data includes an additional 26% based on the mean failure strain of the avian acrocoracohumeral ligament [1]. The ROM values represent the maximal interpolated value from the ROM simulations for the XROMM coordinates. The XROMM values represent the maximal value per ligament calculated across all XROMM trials scaled to each specimen. Nomenclature follows Fürbringer [2]. Abbreviations: LAcH, *Ligamentum (L) acrocoraco-humerale*; LCoHd, *L. coraco-humerale dorsale*; LIcCa, *L. intracapsulare coracoideus anterius*; LScHd, *L. scapulo-humerale dorsale*; LScHl, *L. scapulo-humerale laterale*; LScHp, *L. scapulo-humerale posterius.*

**Supplementary Table S2. Target joint proximity**

| Specimen | Target [mm] | Mean [mm] | Min [mm] | Max [mm] | Overlap |
| --- | --- | --- | --- | --- | --- |
| RLP2 | 1.863 | 1.805 | 0.849 | 2.979 | 0.569 |
| RLP3 | 1.779 | 1.687 | 0.466 | 3.536 | 0.573 |
| RLP4 | 1.752 | 1.877 | 1.191 | 2.969 | 0.409 |
| Average | 1.798 | 1.790 | 0.835 | 3.161 | 0.517 |

The target joint proximity was defined as half the radius of the sphere fitted to the glenoid articular surfaces. The average, minimum and maximum values were determined from the CT scan position. Overlap was defined as the proportion of the articular distances that remained below the target threshold [3].

**Supplementary Information**

***Anatomical and joint coordinate system setup***

A sphere was fitted to the combined glenoid articular surfaces of both coracoid and scapula. The glenoid ACS (Figure S2) was positioned with its origin at the glenoid sphere centroid and was constrained to match the orientation of the body ACS.

An ellipsoid was fitted to the humeral head articular surface instead of a sphere, as the articular surface is elongate and an ellipsoid therefore defines the joint centre more accurately [4], but see below. Due the highly three-dimensional movement of the ulna and radius relative to the humerus in birds [5], a spherical joint rather than a hinge-joint might better approximate their motion. Therefore, spheres were fitted individually to the spherical ulnar and radial condyles, rather than a cylinder across them. The vector from the radial sphere (lateral) to the ulnar sphere (medial) represented the principal vector of the distal humerus, while the vector from the centroid of the humeral head ellipsoid to the midpoint between the distal spheres represented the long-axis vector of the humerus, from which the cardinal axes (*Z*- and *X*-axis, respectively) were derived with their origins at their respective centroids [6] (Figure S2). The other axes of the humerus were derived from the cardinal axes by vector math; that is, crossing the proximal *X-*axis with the distal *Z-*axis and vice versa resulted in the respective *Y-*axes.

The ulna and radius were set up as individual elements and not combined into a single zeugopodial element, unlike previous studies (e.g., Baier *et al.* 2013; Heers *et al.* 2016; Gatesy *et al.* 2022) as their individual motion contributes to the wing movement during flight and folding of the wings [5,9]. Cylinders were fit to their distal articular surfaces and planes to their proximal ones, their centroids defining the relative ACS origin. The fitting of a plane and cylinder results in a pair of T-shaped principal vectors, like in the humerus. These defined the cardinal axes from which the *Z-*axis (flexion-axis in the distal ACS) and *X-*axis (long-axis; in the proximal ACS) were established. The other axes for the ulna and radius ACSs were then derived from these cardinal axes by vector math [6].

For the carpometacarpus, a cylinder was fit to the proximal articular surface delimited by the *trochleae carpalis ventralis et dorsalis*, and a plane was fit to the articular surface for digit II at its distal end. As in the other long bones, the cylinder and plane resulted in T-shaped vectors from which the ACSs were defined. The carpal JCS was defined between the distal ulna ACS and the proximal carpometacarpus ACS.

Unlike the previously discussed ACSs, the body ACS was not defined by individual bones but instead by three landmarks on the skeleton: locators were placed at the median trabecula of the sternum and at the cranialmost point of each coracoid. The origin of the body ACS was then defined as the midpoint of these three locators. The vector from the body ACS origin to the median trabecula and the vector from one coracoid locator to the other formed the T-shaped principal vectors from which the *Y*- and *X*-axes were established. Crossing the *Y-* and *X-*axes resulted in the *Z-*axis, positive pointing upwards.

***Functional joint centre calculation***

The functional centre of the shoulder joint was estimated based upon cadaveric manipulation of the RLP specimens and the resulting XROMM data across all specimens. Individual translational offsets were size-normalised by the minimal diaphyseal circumference of the humerus of each respective specimen [10] and the datasets from all specimens were subsequently combined. The humeral circumference of each µCT scanned specimen was calculated in Autodesk Maya based on a cutting plane at the narrowest part of the humeral shaft using an implementation of Andrew's monotone chain convex hull algorithm [11].

The functional joint centre was assumed to be represented by the mean joint centre position. 4x4 transformation matrices were calculated for the normalized 6-DOF movement from one time step (frame) to the next across the whole dataset, using the *KineMat* toolset for Matlab [12]. Screw axes (i.e., the instantaneous helical axes [13]) of successive frames were calculated from these matrices [14,15]. These screw axes described the motion of a bone relative to another from one frame to the next as a translation along and rotation around a single axis (i.e., a helical motion [13]). The mean joint centre was then determined as the point closest to all finite helical axes in the least squared sense; that is, the point for which the sum of squared distances was minimal [15].

***Comments on mean joint centre***

Due to the substantial sliding (i.e., translational movement) in the RLP shoulder joint neither fitting a sphere to the glenoid articular surface on the coracoid and scapula nor an ellipsoid to the humeral head allowed the shoulder joint centre’s position to be accurately determined (Figure S5). The mean joint centre position, derived from helical axis modelling, was, however, situated within the intersecting area of the two fitted shapes in the μCT scan positions of all specimens. In relation to the glenoid ACS. the mean joint centre was positioned on negative *X* (cranial) and *Y* (lateral), and positive *Z* (dorsal) (Figure S6)*.* In relation to the proximal humeral ACS, the mean joint centre was positioned mostly towards negative *X* (proximal), whereas it had a relatively wide range on the *Y* (coronal) and *Z* axes (mediolateral) (Figure S6). Because the mean joint centre was expressed in the glenoid coordinate system, it remained stationary and always remained within the glenoid sphere. In contrast it was highly dynamic in the humeral coordinate system and sometimes left the humeral ellipsoid. For RLP2 the mean joint centre remained within the humeral ellipsoid in 87.458% of the trial poses, whereas this was the case for 96.726% of the poses in RLP3 and 93.404% in RLP4 (Figure S6).

***Implementation of signed distance field-based simulations in Autodesk Maya***

*ROM simulation approach*

Automated estimation of joint range of motion via optimisation of joint translations was conducted using signed distance fields (SDFs). The scripts optimise contact-based positions across a given set of rotational poses for a set of bone meshes. This is an implementation of the Marai et al. [16] and Lee et al. [17] approach for Autodesk Maya. It creates SDFs for the proximal and distal bone meshes which are then used to calculate intersections between the meshes and the distance between the bones. The (mobile) joint centre position is estimated, and the distal bone mesh is moved into position for a set of prescribed joint orientations. Only feasible joint positions are keyed/exported. The resulting set of frames represents viable joint positions and orientations.

The multiterm objective function $C_{ROM}$ to optimise the translational joint position for each given joint orientation was implemented as follows:

$$C_{ROM}=\underset{\text{joint proximity goal}}{\underbrace{\left( \frac{1}{n_{v}}\sum_{v=1}^{n_{v}} d_{v}-d_{t} \right)^{2}}}+\underset{\text{joint congruency goal}}{\underbrace{\left( \frac{1}{n_{v}}\sum_{v=1}^{n_{v}} \left( d_{v}-\bar{d} \right)^{2} \right)}}$$

subject to the inequality constraint imposed by the SDFs

$$f\left( x_{v},y_{v},z_{v} \right)\geq0$$

where $d_{v}$ is the calculated signed distance for vertex $v$, $n_{v}$ is the number of vertices of the articular surfaces, $d_{t}$ is the target joint proximity, $\bar{d}$ is the mean joint proximity, and $x$, $y$ and $z$ are the Cartesian 3D coordinates of the sample point.

Only joint poses where the average intraarticular distance did not exceed the target joint proximity by more than 7% were deemed viable, as the joints would otherwise have been likely disarticulated. The target joint proximity was determined as half the radius of the sphere fitted to the glenoid articular surfaces (which was a close match to the measured average intra-articular distances from the CT scans, yet does not rely on articulated remains), and represents a joint overlap (*sensu* Bishop et al. [3]) of approximately 0.5 (Supplementary Table S2).

*How to run the ROM simulations*

**Prepare the Maya scene:**

1. Several sets of objects are required:

(1) congruencyMeshes - representing the articular surfaces (e.g., ['prox_art_surf', 'dist_art_surf']).

(2) meshes - representing the bones themselves (e.g., ['prox_mesh', 'dist_mesh']).

(3) fittedShape - representing the fitted shape to the proximal articular surfaces (which is a proxy for the target joint spacing; e.g., 'prox_ftted_sphere').

(4) jointName - representing the joint (e.g., a joint called 'myJoint' if following Manafzadeh and Padian [18]. Note that an attribute called *viable* in jointName is no longer required.

2. Meshes representing the bones (i.e., meshes) need relatively uniform face areas, otherwise large faces might skew vertex normals towards their direction. It is, therefore, important to extrude edges around large faces prior to closing the hole at edges with otherwise acute angles to circumvent this issue (e.g., if the bone meshes have been cut to reduce polycount) prior to executing the Python scripts.

**Execution of the scripts:**

The ROM simulations can be run both from within Maya or directly from the command line/terminal without the need to open the Autodesk Maya application, similar to the AutoBend approach [19]. Download the relevant files from:

<https://github.com/OliverDemuth/MayaSignedDistanceFields/tree/main/ROM>

From within Maya:

1. Copy and paste the *ROMmapper.py* script into the *Python Script Editor* and execute it
2. Copy and paste the *runROMmapper.py* script into the *Python Script Editor*
3. Adjust the user-defined variables in the script and execute it

The optimised translations and the corresponding rotations for each viable pose will be keyed into the translation and rotation attributes of jointName for each viable frame. Only viable poses will be keyed. Maya will become unresponsive until the ROM simulation is done. A progress bar will be updated according to the progress made. The ROM simulations can be cancelled at any time by pressing *Esc* without losing any progress.

From the command line (terminal; Autodesk Maya does not need to be open):

1. Create the following exemplary folder structure and change the directories in the *ROMmapperWrapper.py* script

path = '/your/file/path/ROM/python'

fileDir = '/your/file/path/ROM/maya files'

outDir = '/your/file/path/ROM/results'

2. Copy the following scripts into the python folder according to the *path* specified in the *ROMmapperWrapper.py* script as above

ROMmapperBatch.py

ROMmapperWrapper.py

3. Adjust the user defined variables in *ROMmapperWrapper.py* and save it
4. Execute the script as follows:

For **Windows** in the command prompt execute the following

cd C:\Program Files\Autodesk\Maya<VersionNumber>\bin\

mayapy /your/file/path/ROM/python/ROMmapperWrapper.py

For **macOS** in the terminal execute the following

cd /Applications/Autodesk/maya<VersionNumber>/Maya.app/Contents/bin/

./mayapy /your/file/path/ROM/python/ROMmapperWrapper.py

The optimised translations and the correspondng rotations for each viable pose will be saved as .csv files in the results folder, named according to the respective Maya file (.mb) in the Maya files folder. The ROM simulations cannot be safely aborted and can only be cancelled by closing the command prompt/terminal, however, all progress will be lost.

The Python scripts were written in Python 3 and were tested with Python 3.11 and Autodesk Maya 2025.

*Ligament simulation approach*

Automated estimation of 3D ligament path wrapping around bone meshes was conducted using SDFs. The scripts calculate the shortest distance of a ligament from origin to insertion, wrapping around the bone meshes. This is an implementation of the Marai et al. [20] approach for Autodesk Maya. It creates SDFs for the proximal and distal bone meshes, which are then used to approximate the 3D path of each ligament across them to calculate their shortest path length from origin to insertion while preventing penetration of the bones.

The ligament path was formulated as the following optimisation problem: Find the coordinates of the $n-1$ points between $p_{0}$ and $p_{n}$, so that the Euclidean distance of the path along $p_{0},p_{1},p_{2}, . . .,p_{n}$ is minimal while the distance between each path point and the bony obstacles is non-negative. In the initial guess, the points were equally spaced along the *X*-axis (i.e., their distance was constant, and their $y$ and $z$ values were set to 0). Thus, the length of the shortest path could be approximated by minimising its Euclidean distance only over the $y$ and $z$ coordinates for each point, which we implemented as the following cost function:

$$C_{lig}=\sum_{i=0}^{n-1} \sqrt{{const}^{2}+\left( y_{i+1}-y_{i} \right)^{2}+\left( z_{i+1}-z_{i} \right)^{2}}$$

subject to the inequality constraint imposed by the SDFs

$$f\left( x_{i},y_{i},z_{i} \right)\geq0$$

and the additional inequality constraint to ensure the smoothness of the optimised ligament paths

$$\tan^{-1} \left( \frac{\sqrt{\max_{i} \left\{ \left( y_{i+1}-y_{i} \right)^{2}+\left( z_{i+1}-z_{i} \right)^{2} \right\}}}{const} \right) \leq\frac{\pi}{3}$$

where $x_{i+1}-x_{i}= \frac{1}{n} =const, i=0 . . . n-1.$ This prevents the ligaments from abruptly changing direction or intersecting the bone meshes between two path points.

*How to run the ligament simulations*

**Prepare the Maya scene:**

1. Several sets of objects are required:

(1) jointName - representing the joint (e.g., a joint called 'myJoint' if following Manafzadeh and Padian [18])

(2) meshes - representing the bones (e.g., ['prox_mesh', 'dist_mesh'])

2. Create two locators for each ligament (i.e., for a ligament called *lig** the origin should be named *lig**_orig and the insertion named *lig**_ins) and position them on the meshes accordingly and parent them underneath the respective elements/joints.
3. For each ligament create a float attribute at jointName and name it accordingly. Make sure that the naming convention for each ligament *lig** and the locators representing their origins and insertions are correct (see above). Make sure to remove the *viable* attribute from jointName if previously followed [18] before executing the ligament calculations.
4. Meshes representing the bones (i.e., meshes) need relatively uniform face areas, otherwise large faces might skew vertex normals towards their direction. It is, therefore, important to extrude edges around large faces prior to closing the hole at edges with otherwise acute angles to circumvent this issue (e.g., if the bone meshes have been cut to reduce polycount) prior to executing the Python scripts.

**Execution of the scripts:**

The ROM simulations can be run both from within Maya or directly from the command line/terminal without the need to open the Autodesk Maya application, similar to the AutoBend approach [19]. Download the relevant files from:

<https://github.com/OliverDemuth/MayaSignedDistanceFields/tree/main/ligaments>

From within Maya:

1. Copy and paste the *ligamentCalculation.py* script into the *Python Script Editor* and execute it.
2. Copy and paste the *runLigamentCalculation.py* script into the *Python Script Editor.*
3. Adjust the user-defined variables in the script and execute it.

The ligament lengths will be keyed into the attributes of jointName for each frame. Maya will become unresponsive until the calculations are done. A progress bar will be updated according to the progress made. The ligament length calculations can be cancelled at any time by pressing *Esc* without losing any progress.

From the command line (terminal; Autodesk Maya does not need to be open):

1. Create the following exemplary folder structure and change the directories in the *ligamentCalculationWrapper.py* script

path = '/your/file/path/ligaments/python'

fileDir = '/your/file/path/ligaments/maya files'

outDir = '/your/file/path/ligaments/results'

2. Copy the following scripts into the python folder according to the *path* specified in the *ligamentCalculationWrapper.py* script as above

ligamentCalculationBatch.py

ligamentCalculationWrapper.py

3. Adjust the user-defined variables in *ligamentCalculationWrapper.py* and save it
4. Execute the script as follows:

For **Windows** in the command prompt execute the following

cd C:\Program Files\Autodesk\Maya<VersionNumber>\bin\

mayapy /your/file/path/ligaments/python/ligamentCalculationWrapper.py

For **macOS** in the terminal execute the following

cd /Applications/Autodesk/maya<VersionNumber>/Maya.app/Contents/bin/

./mayapy /your/file/path/ligaments/python/ligamentCalculationWrapper.py

The ligament lengths will be saved as .csv files in the results folder, named according to the respective Maya file (.mb) in the Maya files folder. The ligament length calculations cannot be safely aborted and can only be cancelled by closing the command prompt/terminal, however, all progress will be lost.

The Python scripts were written in Python 3 and were tested with Python 3.11 and Autodesk Maya 2025.
